# UMGAP: the Unipept MetaGenomics Analysis Pipeline

**DOI:** 10.1101/2021.05.18.444604

**Authors:** Felix Van der Jeugt, Rien Maertens, Aranka Steyaert, Pieter Verschaffelt, Caroline De Tender, Peter Dawyndt, Bart Mesuere

## Abstract

Shotgun metagenomics is now commonplace to gain insights into communities from diverse environments, but fast, memory-friendly, and accurate tools are needed for deep taxonomic analysis of the metagenome data. To meet this need we developed UMGAP, a highly versatile open source command line tool implemented in Rust for taxonomic profiling of shotgun metagenomes. It differs from state-of-the-art tools in its use of protein code regions identified in short reads for robust taxonomic identifications, a broad-spectrum index that can identify both archaea, bacteria, eukaryotes and viruses, a non-monolithic design, and support for interactive visualizations of complex biodiversities.

## Introduction

Biodiversity, in many environments, is formed by complex communities of archaea, bacteria, eukaryotes, and viruses. Most of these organisms are hard to isolate and culture in lab conditions, so getting insight into which species are present in these environments and estimating their abundances nowadays routinely relies on metagenomics (Hugenholtz and Tyson 2008): a combination of high-throughput DNA sequencing and computational methods that bypass the cultivation step to enable genomic analysis. In particular, shotgun metagenomics, the non-targeted sequencing of all genomes in an environmental sample, is applied more often (Quince, C. et al. 2017), as it allows the profiling of both taxonomic composition and functional potential of the sample.

In general, computational approaches for taxonomic profiling of metagenomics data from high-complexity environments directly process the reads by either assembling them into larger contigs before profiling (Peng, Y. et al. 2011, 2012; Namiki, T. et al. 2012; Simpson, J.T., Wong, K., and Jackman, S.D. 2009; Boisvert, S. et al. 2012; Pell, J. et al. 2012) or by individually mapping them to DNA sequence databases (Huson, D.H. et al. 2011; Brady, A. and Salzberg, S.L. 2009; Wood, D.E. and Salzberg, S.L. 2014), e.g., compiled from publicly available reference genomes. As the latter approach is carried out without assembly, it can mitigate assembly problems, speed up computations, and enable profiling of low-abundance organisms that cannot be assembled de novo (Quince, C. et al. 2017). Mapping a read onto a reference database usually either applies inexact string matching algorithms on the entire read or breaks it into shorter fragments before applying exact string matching algorithms.

In this paper we explore the alternative route of first translating the protein coding regions in the reads of a shotgun metagenomics data set. We then map the resulting protein fragments onto a reference protein sequence database. Because protein sequences are more conserved than the genes encoding them (Watson, J.D. et al. 2008), this might alleviate the limitation that environmental samples contain strains that are not covered in reference databases or even belong to yet uncharacterized species. In addition, it allows us to adapt the high-performance mapping algorithms based on periodic builds of a UniProtKB-based index (Boeckmann et al. 2003) for mapping tryptic peptides that we developed for shotgun metaproteomics analysis in Unipept (Mesuere, B. et al. 2012, 2015; Gurdeep Singh, R. et al. 2019; Verschaffelt, P. et al. 2020). Mapping against the complete UniProt Knowledgebase has the advantage that it covers all domains of life in a single general-purpose analysis, compared to using one or more reference databases of selected genomes.

The general steps involved in our approach for taxonomic profiling of a DNA read are outlined in Figure 1. After identifying and translating the (partial) protein coding genes in the read, the protein fragments are split into short peptides. Each individual peptide is then mapped onto a precomputed consensus taxon derived from all proteins containing the peptide in the reference database. As a final step we derive a consensus taxon for the read from the consensus taxa of its individual peptides. For paired-end sequencing, the information content in the final step increases after merging the individual peptides from a read pair, as it is guaranteed that both reads originate from the same organism.

**Figure 1:**
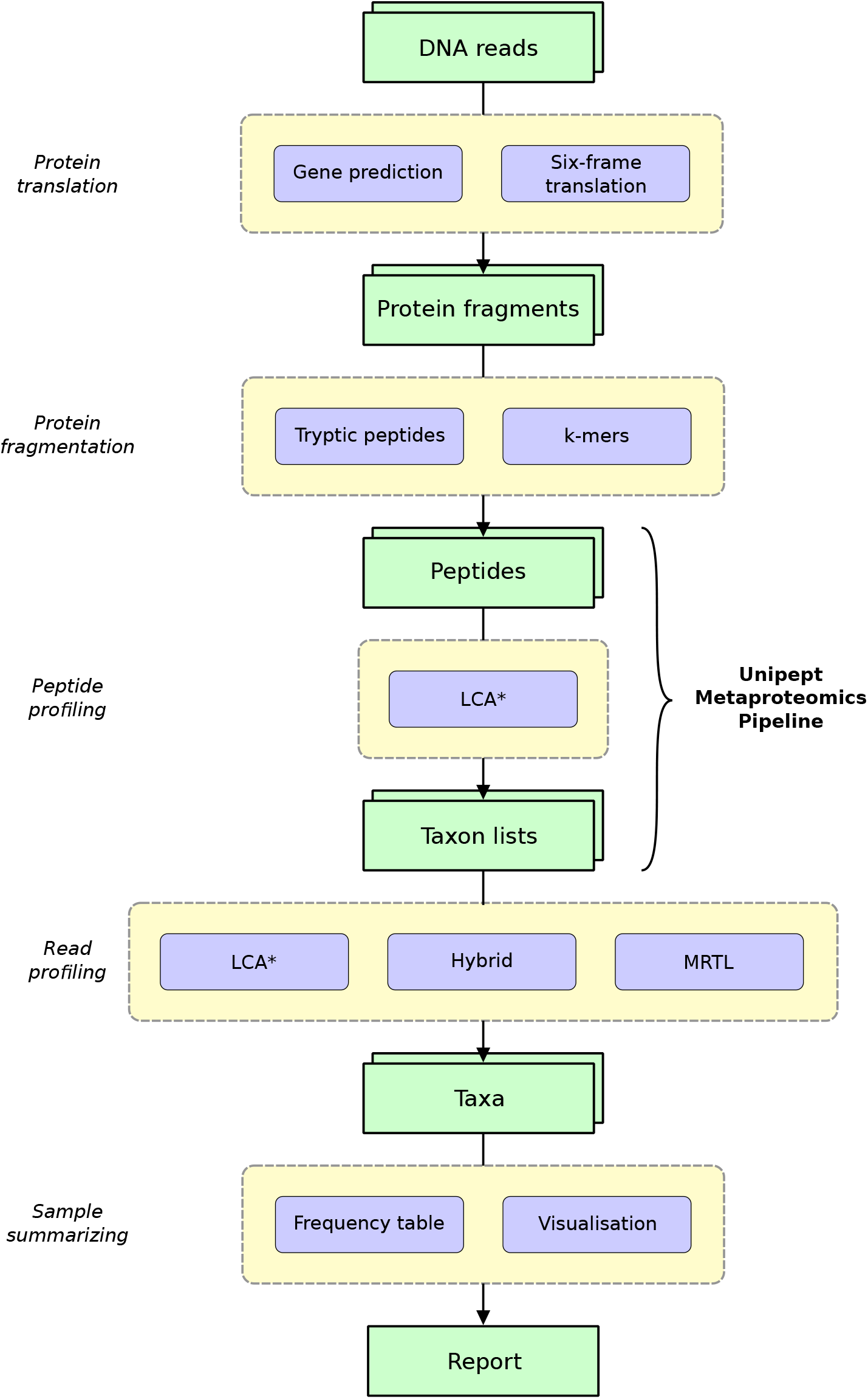
Outline of the Unipept Metagenomics Analysis Pipeline (UMGAP). Green blocks represent data types, purple blocks represent tools. Horizontally aligned purple blocks grouped in yellow boxes are alternative approaches for the same general step of the pipeline.

Each individual step in the above process can be tackled using a multitude of strategies. To explore which strategy performs best and how the combination of alternative strategies leads to different trade-offs with respect to runtime, memory footprint and predictive accuracy, we have implemented the Unipept Metagenomics Analysis Pipeline (UMGAP) according to the Unix philosophy (Raymond 2003). The result is a modular suite of 20 versatile filters (commands that read from standard input and write to standard output) that each implement a single operation and that can be seamlessly combined into a single data processing pipeline. All filters are implemented in [Rust](https://www.rust-lang.org/) and support parallelisation to achieve optimal performance. As some filters implement alternative strategies of the same operation, we have performed a parameter sweep to collect performance metrics of all relevant combinations of alternative strategies. Based on our observations from this parameter sweep, we have selected six preconfigured pipelines with different performance trade-offs whose results have been compared in an established benchmark (Lindgreen, S., Adair, K.L., and Gardner, P.P. 2016) to a selection of state-of-the-art shotgun metagenomics taxonomic profiling tools.

UMGAP has been open sourced on [GitHub](https://github.com/unipept/umgap) under the MIT License. [Documentation](https://unipept.ugent.be/umgap) and [case studies](https://unipept.ugent.be/umgap/casestudies) are available on the Unipept website.

## Methods

UMGAP performs taxonomic profiling of individual reads or read pairs in a shotgun metagenomics data set. Results can be summarized for the entire data set, either as a hierarchical frequency table containing each identified taxon or as an interactive taxonomic visualization. The pipeline executes a multi-step process and provides fast implementations of alternative strategies for every step of the analysis. In this section, we chronologically discuss the successive steps of the generic pipeline, together with their alternative strategies.

### Protein translation

UMGAP does not profile a read directly at the DNA level. Instead, protein fragments translated from the coding regions in the read are matched. Non-coding regions are ignored *a priori* (which might impact the sensitivity compared to identification strategies that use the entire read) and extra steps are required to find coding regions and protein translations (which might negatively impact both performance and accuracy). However, the more conserved nature of proteins compared to DNA might lead to better generalizations when it comes to profiling environmental strains that have no perfect match in the reference database (Watson, J.D. et al. 2008). Two approaches are supported: one based on gene prediction in short reads and one based on a full six-frame translation (Figure 2).

**Figure 2:**
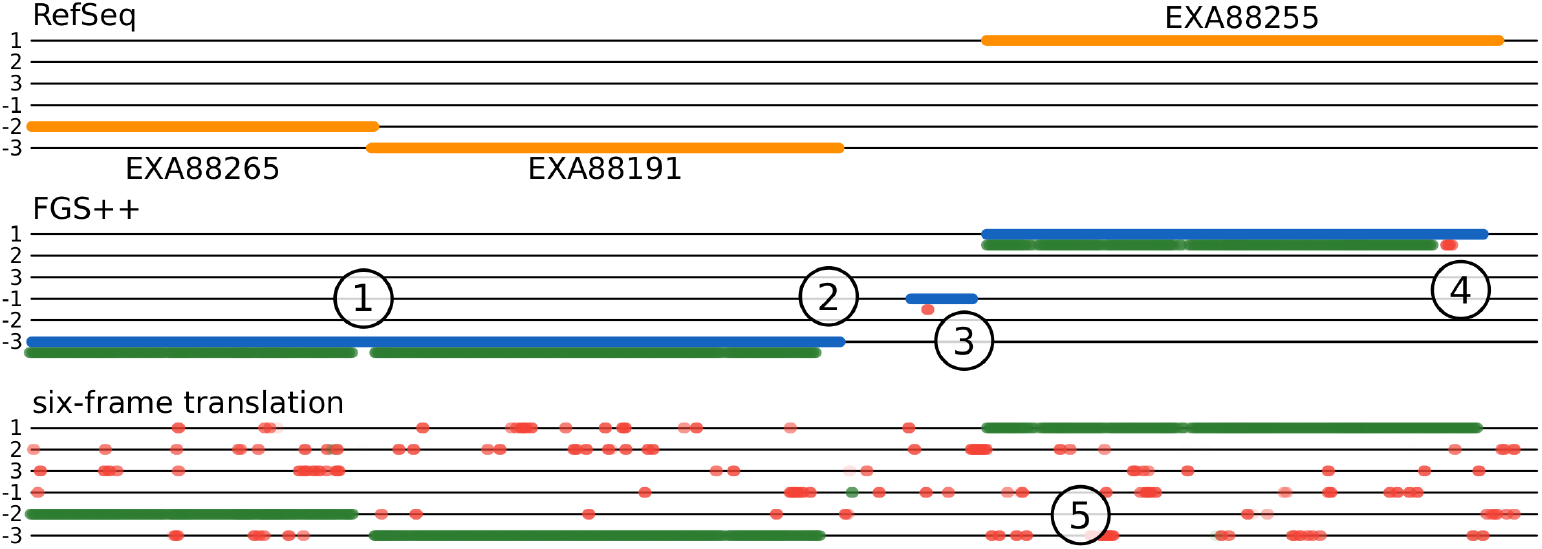
Sample DNA fragment extracted from the *Acinetobacter baumannii* 118362 genome (NCBI Assembly ASM58051v1, positions 37.700-39.530) containing three RefSeq annotated coding regions of a major Facilitator Superfamily protein (EXA88265), a tetR family protein (EXA88191) and a translocator family protein (EXA88255), marked with yellow lines (top). Blue lines indicate coding regions predicted by FGS. Green dots indicate starting positions of 9-mers with an LCA* on the *A. baumannii* lineage (true positive identifications). Red dots indicate starting positions of 9-mers with an LCA* outside the *A. baumannii* lineage (false positive identifications). Opacity of colored dots indicates depth in the taxonomic tree: opaque colors indicate highly specific LCA* (species level) and translucent colors indicate nonspecific LCA*. This example illustrates the following general observations: (1) the frameshift-correcting topology of the FGS hidden Markov model often incorrectly interprets coding regions of genes that are very close or overlapping as frameshifts and glues them together; (2) missing dots at the end of coding regions is merely an artefact of the visualization: the last 8 codons (24 bases) are never starting positions of *k*-mers; (3) FGS may identify false coding regions or (4) frame shifts, but the extracted peptides from those and (5) translations from non-coding regions in a six-frame translation are mostly filtered automatically as they have no exact match with any UniProt protein or can be filtered with additional heuristics.

#### Gene prediction

In theory, UMGAP may use any gene predictor capable of identifying coding regions and their translations in short reads. Not all gene predictors can do this task accurately, as reads might contain only partial coding regions with start and/or stop sections missing. The translation table to be used is also *a priori* unknown.

We used both FragGeneScan (FGS) (Rho, M., Tang, H., and Ye, Y. 2010) and [FragGeneScan++](https://github.com/unipept/FragGeneScanPlusPlus) (FGS++) for testing and benchmarking purposes. FGS has a custom Hidden Markov Model whose topology especially addresses the problem of finding genes in short and error-prone reads, correcting frameshifts resulting from read errors. FragGeneScan-Plus (FGS+) (Kim et al., n.d.) is a faster implementation of FGS. As FGS+ is no longer actively supported, our own implementation with improved multithreading support and several critical bug fixes has been released as FGS++.

FGS and its derivatives are functionally equivalent and can thus be plugged into UMGAP interchangeably. They perform gene prediction relatively fast and accurate. However due to their predictive nature, false negatives and, to a lesser extent, false positives might have a negative impact on downstream steps of the pipeline. Especially missed coding regions may lead to information loss and reduced precision of the pipeline.

#### Six-frame translation

Translation of all coding regions is guaranteed when applied on all six reading frames of an error-free read, but at least 83.33% false positives (5 out of 6 reading frames) need to be filtered away downstream. UMGAP implements this strategy without attempting to correct for read errors at this stage, as they only result in local information loss in downstream steps. The translation table is user-specified, without an attempt to derive it from the data or using multiple tables. While this approach might lead to increased sensitivity compared to gene prediction, it yields at least a sixfold increase in the data volume that needs to be processed in downstream analysis.

### Protein fragmentation

All (partial) proteins that are putatively translated from the read are matched against the complete UniProt knowledgebase (Kim et al., n.d.; Magrane, M. and UniProt Consortium 2011). Direct full-length exact matching is not feasible due to natural variation and read errors. Even though fast heuristics exist for full-length inexact matching or alignment (Altschul et al. 1990), it remains a relatively slow approach. Instead, UMGAP achieves high-performance inexact matching of protein fragments by breaking them down into short peptides. Two approaches are supported: non-overlapping variable-length peptides and overlapping fixed-length peptides.

#### Tryptic peptides

UMGAP breaks protein fragments into non-overlapping variable-length peptides by splitting after each lysine (K) or arginine (R) residue that is not followed by proline (P). This is the classic *in silico* emulation of a trypsin digest, the most widespread protein digestion used for mass spectrometry (Vandermarliere, E., Mueller, M., and Martens, L. 2013). It is a random fragmentation strategy in the context of metagenomics, but finds its roots in the Unipept metaproteomics analysis pipeline as the initial starting point for UMGAP, and is merely an attempt to reuse part of the metaproteomics processing pipeline for metagenomics analysis. Applying this fragmentation strategy to all proteins in the UniProt Knowledgebase (2020/04 release) yields a collection of tryptic peptides with an average length of 17.671 amino acids (with peptides shorter than 5 or longer than 50 discarded).

#### *K*-mers

With overlapping fixed-length peptides or *k*-mers, the only parameter is the length *k* of the peptides. Choosing smaller *k* leads to more spurious hits when matching the *k*-mers of a protein fragment against the *k*-mers inferred from a reference protein database. Choosing larger *k* increases the impact of natural variation and read errors. Because protein fragments are reduced to all their overlapping *k*-mers, the number of resulting peptides increases *k*-fold compared to using tryptic peptides. So, choosing larger *k* also increases the total length of all peptides and thus the memory footprint for indexing them. It also increases the number of lookups that need to be done during peptide profiling. Finding a well-balanced peptide length *k* is therefore crucial.

Because UMGAP uses exact matching for mapping peptides derived from a read onto peptides derived from the proteins in a reference database, read errors and natural variation usually have a lower impact on *k*-mers than on tryptic peptides. This is illustrated in Figure 3 for a single nucleotide polymorphism (either as a read error or a natural variation).

**Figure 3:**
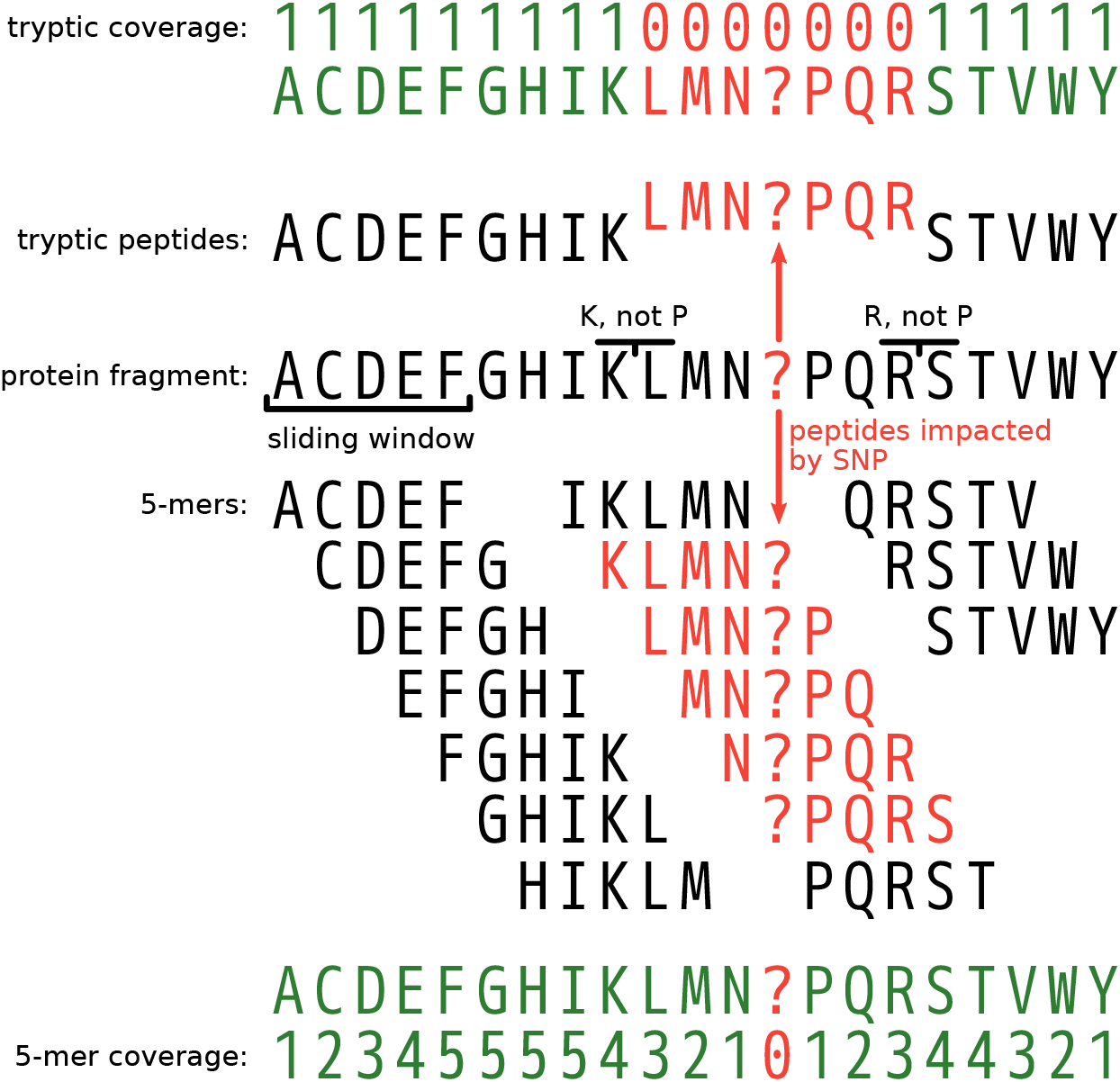
Impact of a single nucleotide polymorphism (SNP) on the peptide coverage of a protein fragment. Because tryptic peptides are non-overlapping, each read position is covered by exactly one tryptic peptide. A SNP modifies a single tryptic peptide, but all positions around the SNP covered by the modified peptide are no longer covered (correctly) after matching (53 read positions on average). Because *k*-mers are overlapping, each read position is covered by k peptides, except at the ends of the protein fragments. A SNP modifies k peptides, but apart from reduced peptide coverage around the SNP, only a single codon is no longer covered (correctly) after matching. The redundancy of the overlapping *k*-mers therefore makes up for a reduced impact of SNPs compared to non-overlapping tryptic peptides, at the cost of larger data volumes that need to be processed.

### Peptide profiling

Fragmentation of all partial proteins found in a read yields a list of peptides (tryptic peptides or *k*-mers). Each peptide may have an associated consensus taxon that is looked up in an index structure. Overall, these lookups are the most time-consuming step in the pipeline, so performance is of utmost importance.

Upon each UniProt release, the Unipept team builds and publishes new indexes from tryptic peptides and 9-mers extracted from all UniProt proteins in the knowledgebase (available online). Each of these peptides is associated with the LCA* consensus taxon computed from the set of taxonomic annotations on all Uniprot proteins that exactly match the peptide (Mesuere, B. et al. 2012). LCA* is the most specific taxon that does not contradict any taxon in the set, i.e., all taxa in the set must either be descendants or ancestors of the LCA* in the NCBI Taxonomy (Federhen 2012). See the read profiling step for a detailed discussion on the LCA* algorithm introduced by UMGAP as a variation on the lowest common ancestor (LCA) algorithm.

UMGAP uses a finite state transducer (Gallant n.d.) as its index structure to lookup the LCA* consensus taxon for each peptide extracted from a read. This index structure supports high-performance and parallel lookups, supports both fixed and variable length peptides, and has a relatively small memory footprint. The latter is important, given the large number of peptides extracted from UniProt. The index should be loaded in process memory, but UMGAP can also operate with an on-disk index and very little memory at the cost of performance.

The FST maps each peptide extracted from a UniProt protein to the NCBI Taxonomy Identifier (an integer) of the LCA* associated with the peptide. It is a flow graph whose edges are labeled with amino acids and integers. Peptides are matched by following the path of their amino acid sequence. The sum of the integers along this path corresponds the identifier of the LCA*. Where tries are ordered tree data structures that take advantage of common prefixes to reduce the memory footprint, FSTs are even more compressed by taking both common prefixes and suffixes into account (Supplementary Figure 10).

For UniProt release 2020-04, a 19.3 GiB FST-index maps all 1.2 billion tryptic peptides to their LCA* and a 132.9 GiB FST-index maps all 17 billion 9-mers to their LCA*. We also experimented with other k-mer lengths, but precision dropped significantly for *k* ≤ 7 (Figure 4) and the index size became too large for *k* ≥ 10. The only viable options were *k* = 8 and *k* = 9, with the latter giving the best balance between index size and accuracy of read profiling.

**Figure 4:**
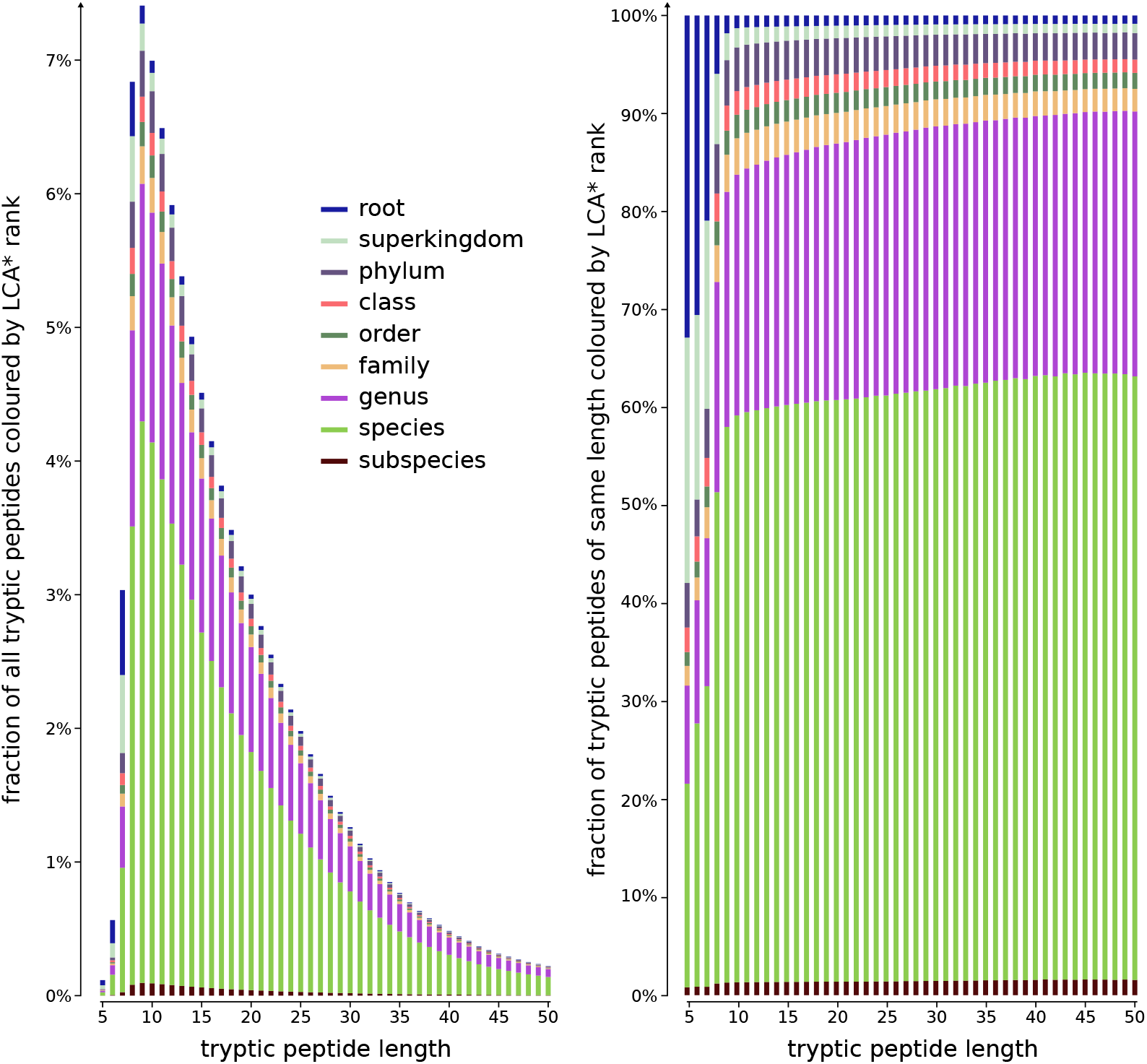
All tryptic peptides found in the UniprotKB (release 2020-04-22), classified by length and associated LCA* rank. Fractions of peptides (y-axis) across all peptide lengths (left) or per peptide length (right). Short tryptic peptides are more frequently associated with less specific ranks in the NCBI Taxonomy and therefore have a lower information content. Relative taxonomic information content (depth of LCA* rank in tree of life) is low for short peptides (length 8 and below). Because tryptic peptides of length *k* are a random sample of all *k*-mers, similar ratios and conclusions are expected should this analysis be repeated for all *k*-mers in UniprotKB across all lengths *k*.

### Peptide filtering

Protein fragmentation may yield false positives: peptides that do not occur in proteins encoded in the read. Most false positives are automatically filtered as they have no exact match with any UniProt protein. As a result, they cannot be associated with a taxon during peptide profiling. This is the case for most peptides from translations of wrong gene predictions or outside coding regions in a six-frame translation (Figure 2). But peptide profiling itself may also yield false positive identifications: peptides associated with an inconsistent taxon, i.e., a taxon that is not the correct taxon or one of its ancestors in the NCBI Taxonomy tree. This could be the case for both true and false positive peptides from protein fragmentation. Peptide filtering aims at strongly reducing the number of false positive identifications, while keeping most true positives. UMGAP supports three kinds of filters.

#### Short tryptic peptides

Analysis on UniProt proteins shows that short tryptic peptides are typically associated with highly unspecific LCA* consensus taxa, i.e., taxa at or close to the root of the NCBI Taxonomy tree (Mesuere, B. et al. 2012) (Figure 4). Because these peptides match proteins occurring across all domains of life, they do not provide a strong taxonomic signal that could be useful in downstream steps of the pipeline. In addition, by being short they often cause spurious matches during peptide profiling. UMGAP can skip very short tryptic peptides, e.g., having less than 6 amino acids.

#### Low-frequency identifications

In the context of peptide profiling, true positive identifications should come from the same lineage in the NCBI Taxonomy tree, whereas false positives are randomly scattered across the tree. Since one read typically yields many peptides that may each have an associated taxon, identifications along the correct lineage are expected to occur with high frequency and false positives are expected to occur with low frequency. Therefore UMGAP can skip peptides associated with low-frequency identifications.

#### Seed-and-extend

The (partial) proteins in the read are typically fragmented into multiple peptides and it is expected that neighboring peptides have similar identifications (Figure 2). It therefore seems natural to use a seed-and-extend approach to exploit this expected local conservation of identifications. Peptides are first scanned to find seeds: *s* or more successive peptides that are associated to the same LCA*. With increased minimum seed size *s*, the precision of the pipeline will increase, and its sensitivity will decrease. Each seed is then extended in both directions to neighboring seeds and individual peptides that are bridged by gaps (successive peptides with no associated LCA*) of at most *g* peptides. With increased maximum gap size *g*, the precision of the pipeline will decrease, and its sensitivity will increase. UMGAP can skip peptides that are excluded from extended seeds (Figure 5).

**Figure 5:**
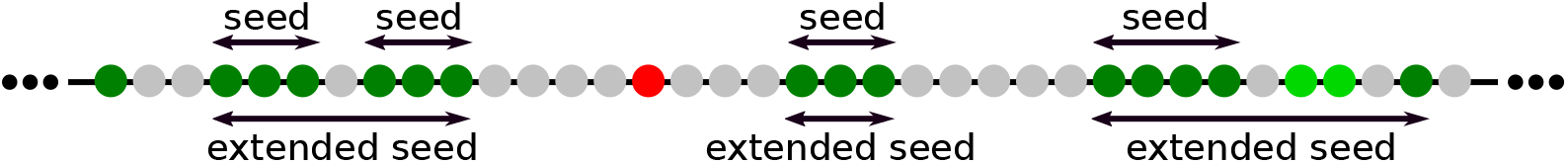
Seed-and-extend strategy for filtering false positive identifications after peptide profiling, with minimum seed size *s* = 3 and maximum gap size *g* = 1. Successive peptides fragmented from (partial) protein are shown as a sequence of dots. Green dots indicate correct identifications (true positives). Red dots indicate wrong identifications (false positives). Brightness of colored dots indicates depth in the taxonomic tree: dark colors indicate highly specific LCA* (species level) and light colors indicate nonspecific LCA*. Grey dots indicate peptides without an associated LCA*.

### Read profiling

Previous steps of the pipeline result in a list of taxonomic identifications, derived from a (filtered) list of peptides extracted from the read. As the read comes from a single organism, it is natural to aggregate these individual identifications that rely on partial data into one global consensus identification. UMGAP supports three heuristics that infer a consensus taxon after mapping a frequency table of the individual identifications onto the NCBI Taxonomy tree (Figure 6). They balance between providing a consensus taxon that is as far away from the root as possible and that allows for a good generalization. The first goal is progressive and leads to a very specific consensus but has to avoid overfitting. The second goal is conservative and takes into account the possibility of false positives among the individual identifications but has to avoid underfitting. All heuristics are implemented with two different data structures: a tree and a range minimum query (Fischer, J. and Heun, V. 2011). Both implementations are functionally equivalent, but the latter gives a faster implementation of the MRTL heuristic because querying ancestry is supported in constant time.

**Figure 6:**
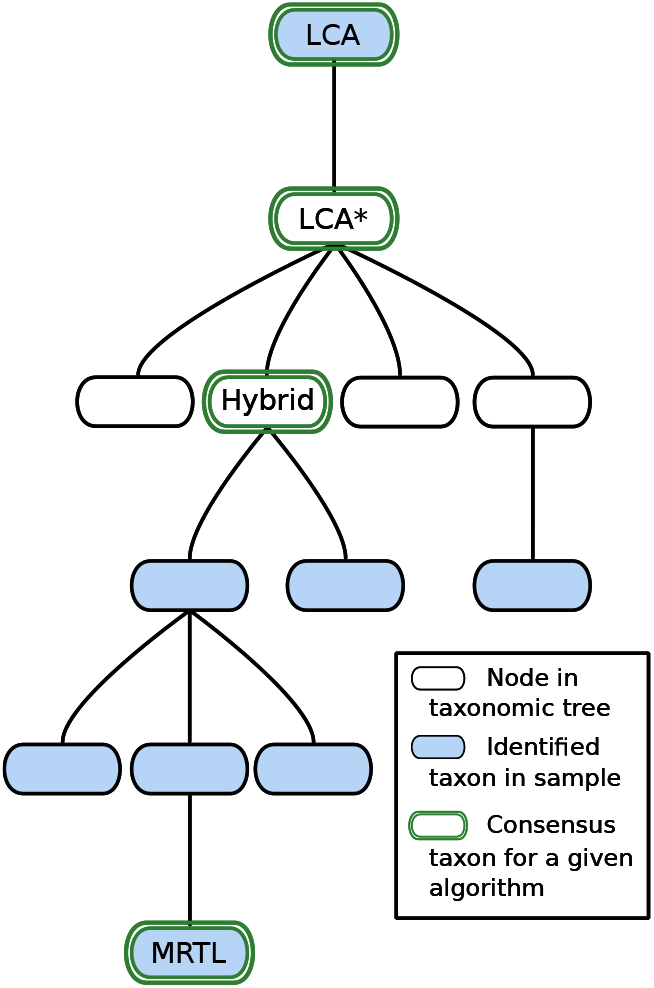
Lowest common ancestor (LCA), LCA*, hybrid_*f*_ and maximum root-to-leaf (MRTL) are four heuristics that determine a consensus taxon for a given list of taxa. All heuristics map the list of taxa onto the NCBI Taxonomy tree, that guides the heuristic towards a consensus taxon. MRTL is the most progressive heuristic and LCA* is the most conservative heuristic, so the MRTL consensus will be deeper in the tree than the LCA* consensus. The hybrid_*f*_ heuristic has a parameter *f* that can be tweaked to yield a result that is either more towards the LCA* consensus or the MRTL consensus.

#### MRTL

Maximum root-to-leaf (Wood, D.E. and Salzberg, S.L. 2014) is the most progressive heuristic. It identifies the consensus among a list of taxa as a taxon having the maximal number of ancestors in the list. Ties are broken arbitrarily. By definition, the consensus taxon is always included in the original list of taxa. This property does not hold for the other two heuristics, and shows that this heuristic might not necessarily be good at generalizing.

#### LCA*

is the most conservative heuristic, though less conservative than a standard lowest common ancestor (LCA). For a given list of taxa, it identifies the consensus taxon as the most specific taxon in the tree that is either an ancestor or a descendant of each taxon in the list. This is the LCA of all taxa in the list, after we have first discarded all ancestors of at least one other taxon in the list. The latter is a measure against underfitting. Because the individual identifications are only based on partial data, it is expected that some identifications might be more specific than others. The LCA* heuristic is also used to compute the consensus taxon during peptide profiling.

#### Hybrid_*f*_

This heuristic has a parameter *f* ∈ [0, 1] that allows to balance between being conservative or progressive: with *f* = 1 this heuristic is the same as LCA* and with *f* = 0 this heuristic is very close to MRTL (the same for most lists of taxa). LCA* can be implemented by starting at the root of the tree and repeatedly descending to the child node whose subtree contains all taxa in the list, until such a child no longer exists (i.e., the taxa in the list are distributed over multiple subtrees). In the latter case, the hybrid heuristic continues descending to the child node whose subtree contains most taxa from the list (with ties broken arbitrarily) if the fraction of the number of descendants in the child node over the number of descendants in the current node is larger than or equal to *f*.

### Summary and visualization

Previous steps assign a consensus taxon to each read (pair). The final step of the pipeline computes a frequency table of all identifications across the entire data set, with the option to filter low-frequency identifications. Another option is to report the frequency table at a user-specified taxonomic rank. Frequency tables are exported in CSV-format, enabling easy postprocessing.

To gain insight into environmental samples with a complex biodiversity, UMGAP also supports rendering taxonomic frequency tables as interactive visualizations (Figure 7) that are automatically made available on a dedicated website. The online service hosting the visualizations also support shareable links (e.g. https://bl.ocks.org/5960ffd859fb17439d7975896f513bc3).

**Figure 7:**
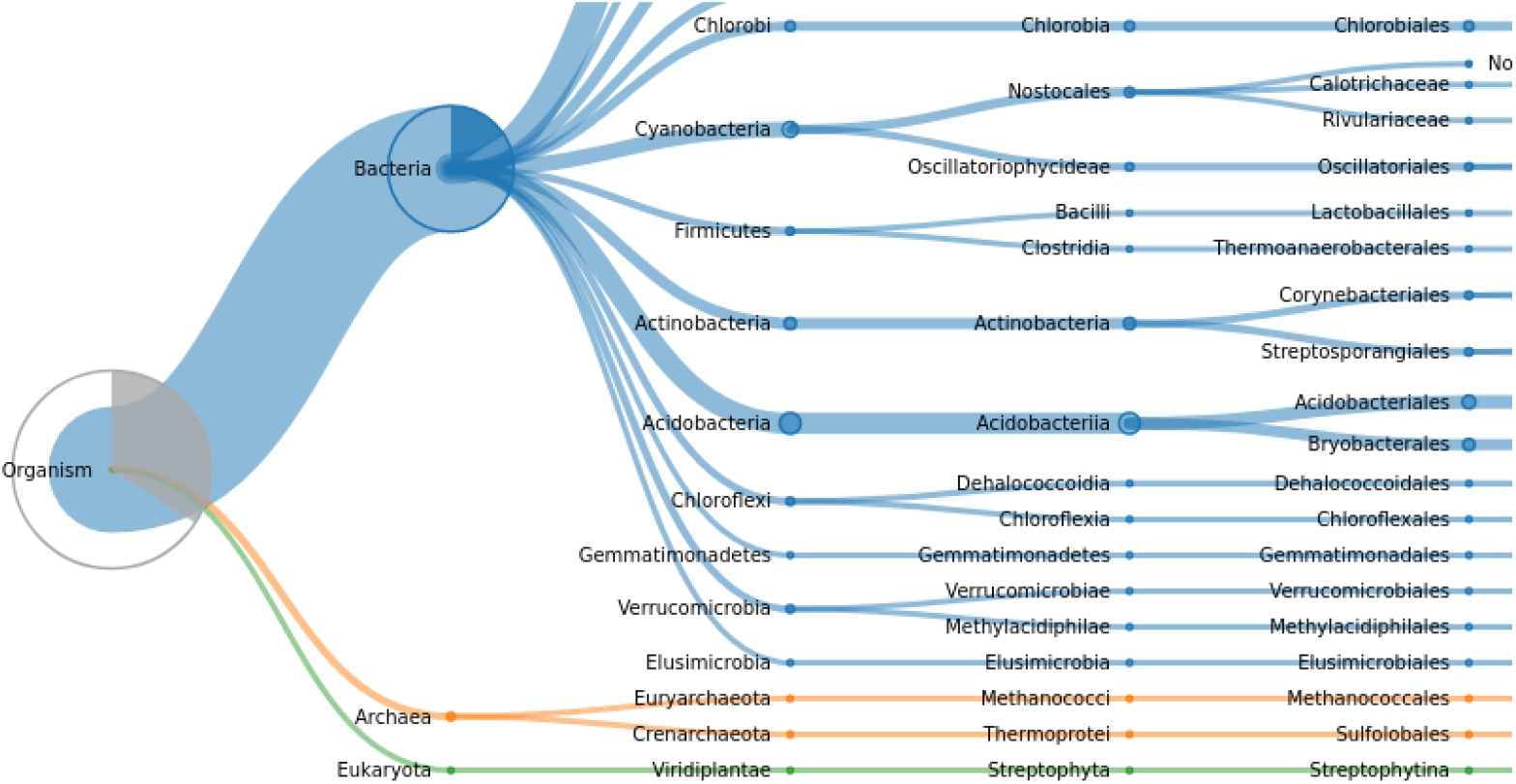
Taxonomic profiling by UMGAP as visualized by the Unipept Web API.

## Results

UMGAP implements multiple strategies for each step in the pipeline (Figure 1), with some strategies also driven by user-specified parameters. Runtime, memory footprint and accuracy of UMGAP were benchmarked as a two-step process. Using some smaller data sets, we first measured and analysed performance metrics for a large number of relevant combinations of strategies and parameter settings. This broad exploration allowed us to investigate how different strategies/parameter settings led to different performance trade-offs. As a result, we defined six preconfigured pipelines with different performance trade-offs. Performance of these configurations has then been compared to a selection of state-of-the shotgun metagenomics taxonomic profiling tools in an established benchmark (Lindgreen, S., Adair, K.L., and Gardner, P.P. 2016) that uses larger data sets.

Both the parameter sweep and the benchmark were executed on a 2.60GHz 16 core Intel® Xeon® CPU E5-2650 v2 CPU with 195GB RAM running Debian 9.8 (stretch).

### Parameter tuning

For protein translation we either used gene prediction or six-frame translation. Both FGS and FGS++ were used for gene prediction, as initial tests had shown slight deviations in predictions generated by the two implementations. In the protein fragmentation step we either used non-overlapping tryptic peptides or overlapping 9-mers. Tryptic peptides were filtered by length, with minimum length ranging from 5 to 10 amino acids and maximum length ranging from 45 to 50 amino acids. Peptide profiling was invariably done using LCA*. Low-frequency identifications were filtered with a minimum number of taxon hits per read that varied between 1 and 5, with a minimum of 1 hit effectively corresponding to no low-frequency identification filtering. For 9-mers, identifications were optionally also filtered using the seed-and-extend strategy with seed size *s* between 2 and 4, and gap size *g* between 0 to 4. Read profiling was done using either MRTL, LCA* or hybrid_*f*_, with parameter *f* either set to 0.25, 0.5 or 0.75.

All variation included in this parameter sweep resulted in 3900 different UMGAP configurations whose performance was evaluated for taxonomic profiling of two metagenome data sets simulated by Wood and Salzberg (Wood, D.E. and Salzberg, S.L. 2014). These data sets are referenced as the HiSeq metagenome and the MiSeq metagenome after the Illumina sequencing platforms whose read error models have been used for simulation. For each data set 1000 reads were simulated from 10 bacterial genomes, for a total of 10.000 reads per data set.

Accuracy of each UMGAP configuration was evaluated at the genus level by computing precision and sensitivity of the taxonomic profiling for each data set. Using the UMGAP snaptaxon tool and guided by the NCBI Taxonomy tree, more specific UMGAP predictions were mapped to the genus level because expected predictions were only known at the genus level for these data sets. True positives (TP) are reads assigned to the expected genus. False positives (FP) are reads assigned to a genus other than the expected genus. False negatives (FN) are reads that UMGAP could not assign at or below the genus level. As these data sets contain no invalid reads, there are no true negatives (TN).

Figure 8 shows the precision and sensitivity of all 3900 UMGAP configurations tested. As expected, the protein fragmentation method has a major influence on sensitivity. The difference in precision is less pronounced at first glance. In general, 9-mer configurations (orange) have a higher sensitivity than tryptic configurations (blue), but they also have a higher runtime and memory footprint. To simplify further discussion, we will treat tryptic and 9-mer configurations separately in what follows.

**Figure 8:**
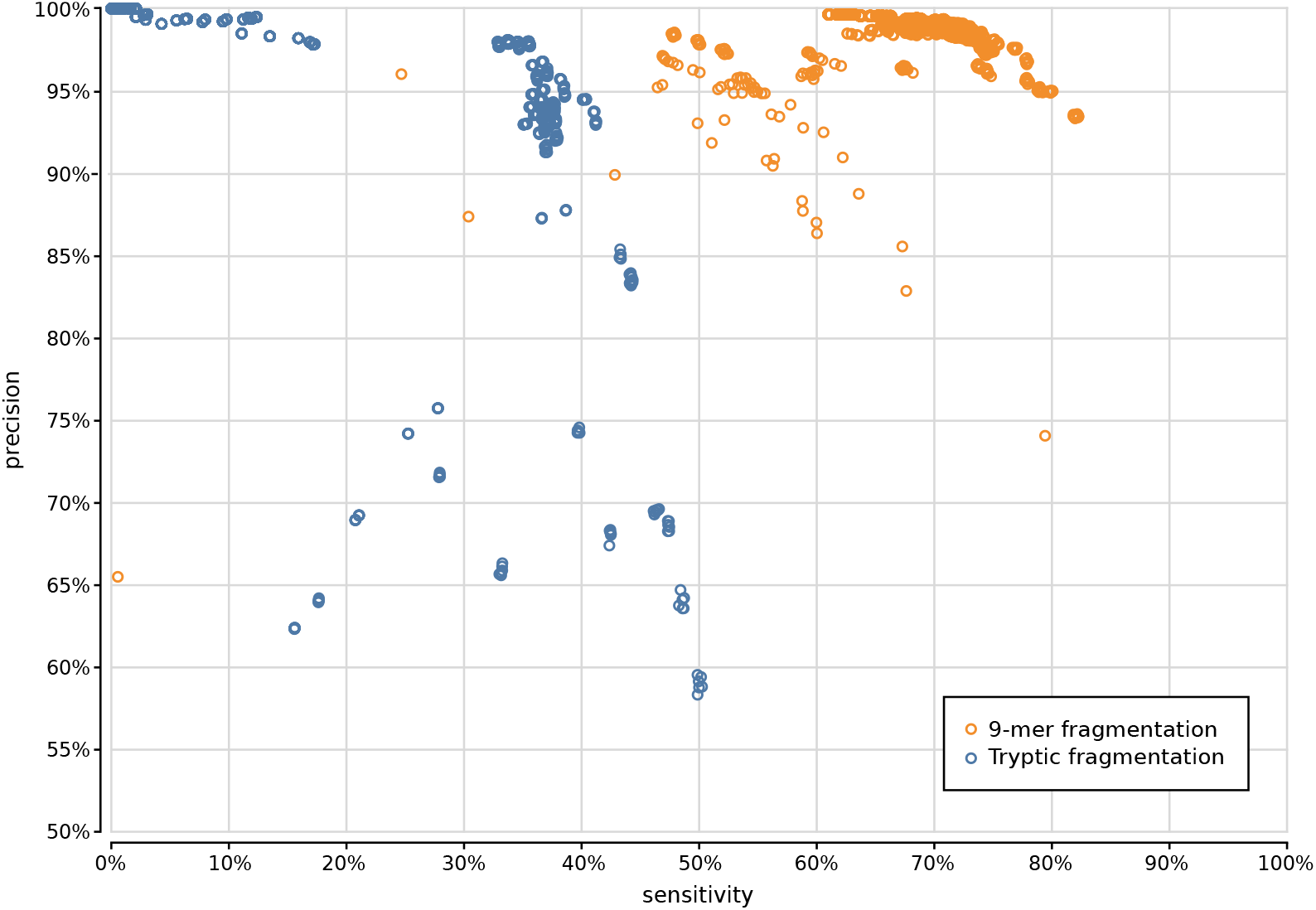
Precision and sensitivity of 3900 UMGAP configurations, with tryptic peptide configurations marked in blue and 9-mer configurations marked in orange.

### Tryptic configurations

If we look at the impact of protein translation on the accuracy of all 2700 tryptic configurations (Supplementary Figure 11), the sensitivity obtained with FGS is slightly better than with FGS++. This may hint on implementation issues with FGS+ (a parallel implementation of FGS) or downstream improvements on FGS after forking FGS+. FGS+ and FGS++ are however faster than FGS. Six-frame translation marginally improves the sensitivity of the tryptic configurations, at the cost of a steep drop in precision if spurious identifications resulting from incorrect translations are not properly filtered after peptide profiling. As it also yields much more work during the peptide profiling step, combining six-frame translation with tryptic peptides proves less favorable.

Shorter peptides have a higher probability of random hits in a protein database. With tryptic peptides, it is therefore recommended to discard very short peptides. In general, we advise to only retain tryptic peptides with a length of at least 9 amino acids (Supplementary Figure 12). We have also investigated the impact of a maximum peptide length cutoff on the accuracy of the predictions, but the effect is negligible except for a marginal gain in the speed of the pipeline.

Tryptic configurations effectively profile only a limited number of peptides per read, such that filtering taxa after peptide profiling must be done carefully to avoid losing valuable information. Discarding taxa that have only been assigned to a single peptide guarantees high precision at the cost of a steep drop in sensitivity (Supplementary Figure 13). This shows that tryptic configurations often profile reads based on a single peptide, increasing the risk of spurious predictions.

The choice of read profiling method has no significant impact on the performance of the pipelines, again because of the limited number of (reliable) peptides per read whose predictions need to be aggregated.

Based on the above observations we have selected two tryptic configurations with good accuracy trade-offs, either favoring higher precision or higher sensitivity:

- **tryptic precision** FGS++, minimum peptide length 5, maximum peptide length 45, minimum 2 taxon hits, MRTL
- **tryptic sensitivity** FGS++, minimum peptide length 9, maximum peptide length 45, no rare taxon filtering, MRTL

### 9-mer configurations

When evaluating all 1200 UMGAP configurations that use 9-mer peptide fragmentation, the first observation is that seed-and-extend filtering has a positive effect on both precision and sensitivity (Supplementary Figure 14). This filtering technique is not useful when working with tryptic peptides, but proves to be highly effective for discarding unreliable identifications after peptide profiling when working with 9-mers. As a result, we recommend to always apply seed-and-extend filtering in 9-mer configurations, and we will only focus on these configurations in any further analysis.

With respect to protein translation method, the same observations concerning accuracy hold as with the tryptic configurations (Supplementary Figure 15). Sensitivity obtained with FGS is slightly better than with FGS++. The sensitivity gain that can be obtained with six-frame translation is more pronounced than with the tryptic configurations, which may make up for the extra work during the peptide profiling step. However, effective filtering of spurious identifications after peptide profiling is still needed in order to avoid poor precision.

Gene prediction is best combined with minimum seed size *s* = 2 for optimal sensitivity and with minimum seed size *s* = 3 for the best trade-off between precision and sensitivity (Supplementary Figure 16). In combination with six-frame translation, better trade-offs between precision and sensitivity are achieved with higher minimum seed size *s*. With gene prediction the low-frequency identifications filter has a higher impact than the chosen read profiling method, whereas the opposite is true for six-frame translation (Supplementary Figures 17, 18, 19, 20). In both cases, the maximum gap size *g* has no significant impact on the accuracy (data not shown).

Based on the above observations we have selected four 9-mer configurations that represent different accuracy trade-offs. Ranging in optimization from precision to sensitivity they use the following UMGAP configurations:

- **max precision** FGS++, minimum 5 taxon hits, seed-and-extend with *s* = 2 and *g* = 2, hybrid_*f*_ with *f* = 0.75
- **high precision** six-frame translation, minimum 4 taxon hits, seed-and-extend with *s* = 3 and *g* = 4, hybrid_*f*_ with *f* = 0.5
- **high sensitivity** six-frame translation, no filtering on low-frequency identifications, seed-and-extend with *s* = 3 and *g* = 0, MRTL
- **max sensitivity** six-frame translation, no filtering on low-frequency identifications, seed-and-extend with *s* = 2 and *g* = 0, MRTL

### Benchmark

The six preconfigured UMGAP pipelines selected from the parameter sweep analysis were compared with the two best-performing shotgun metagenomics analysis tools found in the MetaBenchmark study (Lindgreen, S., Adair, K.L., and Gardner, P.P. 2016) and with the popular Kaiju tool (Menzel, P., Ng, K.L., and Krogh, A. 2016) that was published shortly after the initial benchmark. Kraken (Wood, D.E. and Salzberg, S.L. 2014) and the newer Kraken 2 (Wood, D.E., Lu, J., and Langmead, B. 2019) were run with their default (preloaded) indexes and 16 threads. CLARK (Ounit, R. et al. 2015) was run with 20-mer indexes in full-mode. Because CLARK only supports identifications for a predefined taxonomic rank, we used indexes built from bacterial databases for the taxonomic ranks of phylum, genus, and species. Kaiju was run with its default index.

Our benchmark uses the same experimental setup as the MetaBenchmark study, including its use of two simulated metagenomes that differ in relative abundance of the individual phyla and that have three replicates each. The six data sets contain between 27 and 37 million read pairs simulated from both real, simulated, and shuffled genomes, with read length 100 and mean insert size 500 (standard deviation 25). All data sets contain 20% reads simulated from shuffled genomes that serve as a negative control and also contain reads simulated from genomes that were artificially diverged from a *Leptospira interrogans* reference genome to test the robustness of the tools against natural variation.

In addition to evaluating the accuracy of taxonomic profiling tools at the phylum and genus levels, we also evaluated their accuracy at the species level (Table 1). Using the UMGAP snaptaxon tool and guided by the NCBI Taxonomy tree, predictions more specific than the taxonomic rank under evaluation were mapped to the taxonomic rank under evaluation. Predictions less specific than the taxonomic rank under evaluation were considered as no assignment to any taxon. Reads whose expected identification is less specific than the taxonomic rank under evaluation are ignored. True positives (TP) are non-shuffled reads assigned to the expected taxon. False positives (FP) are non-shuffled reads assigned to a taxon that differs from the expected taxon or shuffled reads assigned to a taxon. True negatives (TN) are shuffled reads not assigned to any taxon. False negatives (FN) are non-shuffled reads not assigned to any taxon.

**Table 1:**
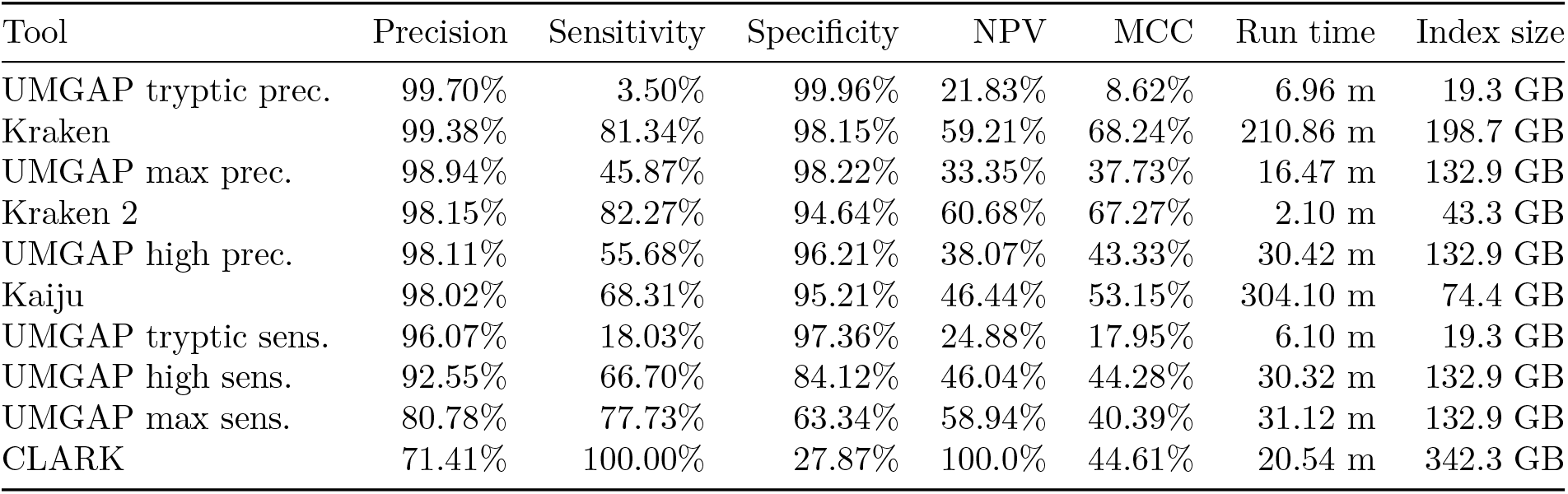
MetaBenchmark performance metrics for ten metagenomics analysis tools sorted by precision. Average numbers for the six simulated data sets are given. Accuracy evaluated at the species level and reported as sensitivity, specificity, precision (positive predictive value), negative predictive value (NPV) and Matthew’s Correlation Coefficient (MCC). Index size reported for CLARK is the sum of the phylum (46.6GiB), genus (149.5GiB) and species (146.3GiB) indexes. Performance metrics at genus and phylum ranks can be found in Supplementary Table 2 and Supplementary Table 3.

| Tool | Precision | Sensitivity | Specificity | NPV | MCC | Run time | Index size |
| --- | --- | --- | --- | --- | --- | --- | --- |
| UMGAP tryptic prec. | 99.70% | 3.50% | 99.96% | 21.83% | 8.62% | 6.96 m | 19.3 GB |
| Kraken | 99.38% | 81.34% | 98.15% | 59.21% | 68.24% | 210.86 m | 198.7 GB |
| UMGAP max prec. | 98.94% | 45.87% | 98.22% | 33.35% | 37.73% | 16.47 m | 132.9 GB |
| Kraken 2 | 98.15% | 82.27% | 94.64% | 60.68% | 67.27% | 2.10 m | 43.3 GB |
| UMGAP high prec. | 98.11% | 55.68% | 96.21% | 38.07% | 43.33% | 30.42 m | 132.9 GB |
| Kaiju | 98.02% | 68.31% | 95.21% | 46.44% | 53.15% | 304.10 m | 74.4 GB |
| UMGAP tryptic sens. | 96.07% | 18.03% | 97.36% | 24.88% | 17.95% | 6.10 m | 19.3 GB |
| UMGAP high sens. | 92.55% | 66.70% | 84.12% | 46.04% | 44.28% | 30.32 m | 132.9 GB |
| UMGAP max sens. | 80.78% | 77.73% | 63.34% | 58.94% | 40.39% | 31.12 m | 132.9 GB |
| CLARK | 71.41% | 100.00% | 27.87% | 100.0% | 44.61% | 20.54 m | 342.3 GB |

In terms of precision the UMGAP tryptic precision configuration marginally surpasses Kraken, with the UMGAP max/high precision configurations, Kraken 2, and Kaiju also showing very high precision rates (Figure 9, Table 1). As expected, the UMGAP pipelines have a lower sensitivity than the other metagenomics pipelines because *a priori* no taxa are assigned to reads that have no or only short overlap with protein coding regions. This benchmark again underscores the difference in sensitivity between the tryptic and 9-mer configurations of UMGAP. Also take into account that precision is a more important accuracy metric than sensitivity for most biological applications, especially with deeply sequenced samples. In terms of speed Kraken 2 is the best-performing tool, with UMGAP’s tryptic configurations following in close range. Clark and the UMGAP 9-mer configurations are still considerably faster than Kraken and Kaiju.

**Figure 9:**
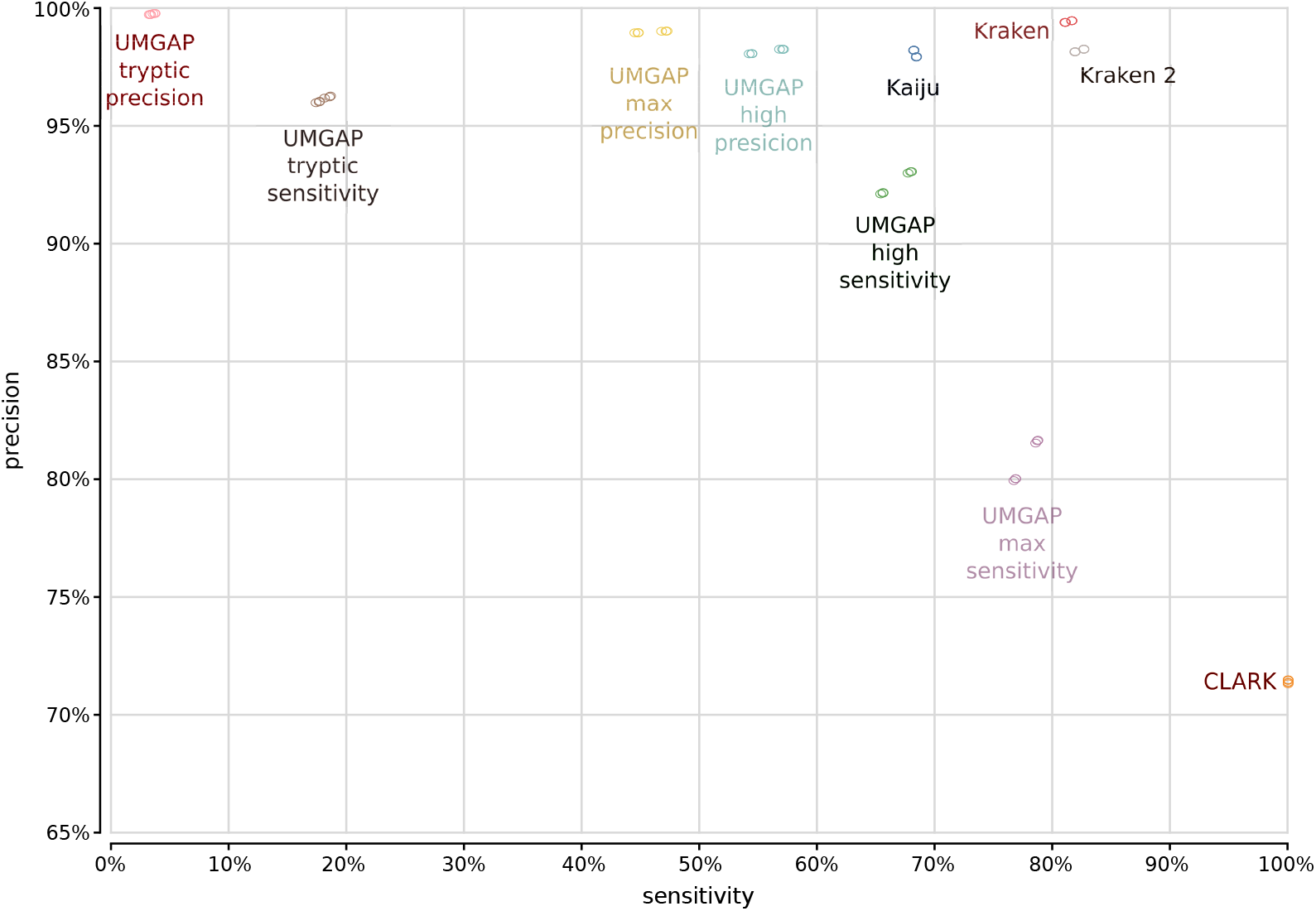
Precision and sensitivity evaluated at the species level for ten metagenomics analysis tools and the two simulated metagenomes of the MetaBenchmark. Dots indicating the accuracy metrics for the three replicates of each simulated metagenome are on top of each other, since each replicate was generated for identical proportions of phyla.

## Discussion

The protein space detour for taxonomic profiling of shotgun metagenomic data sets shows to be very promising. Despite our design choices of an extra protein translation step, a broad spectrum index that can identify both archaea, bacteria, eukaryotes and viruses, and a highly configurable non-monolithic design, UMGAP achieves low runtime, manageable memory footprint and high accuracy that makes it highly competitive with state-of-the-art shotgun metagenomics tools. Integrating the command line tool with the interactive Unipept visualizations (Verschaffelt, P. et al. 2020) also allows exploration and comparison of complex communities.

As a next step, we want to further explore how the protein translation detour can be used to infer the functional capacity of an environmental sample from its metagenome, which is more challenging than inferring biodiversity. Again, Unipept’s function analysis pipeline for metaproteomes could be used as a potential starting point. In addition, both the biodiversity and the functional capacity of a sample could also be derived from its metatranscriptome, which could be analysed using pipelines similar to UMGAP but with an adjusted protein translation step.

## Acknowledgements

We thank Stijn Seghers for his contributions in implementing and benchmarking the initial tryptic peptide components of UMGAP. We thank Niels De Graef for his contributions in implementing and benchmarking the initial prototypes of UMGAP.

## Data

The parameter tuning dataset can be found as the accuracy dataset at [the Kraken website](https://ccb.jhu.edu/software/kraken/). The benchmark dataset can be found at [UC Bioinformatics](http://www.ucbioinformatics.org/metabenchmark.html) The latter are also rehosted at Unipept (A1_1, A1_2, A2_1, A2_2, A3_1, A3_2, B1_1, B1_2, B2_1, B2_2, B3_1, and B3_2) to ensure availability. The created index files are dated by UniprotKB release day and are found at https://unipept.ugent.be/system/umgap/recent/ninemer.fst, https://unipept.ugent.be/system/umgap/recent/tryptic.fst and https://unipept.ugent.be/system/umgap/recent/taxons.tsv.

## Supplementary Tables and Figures

**Table 2:**
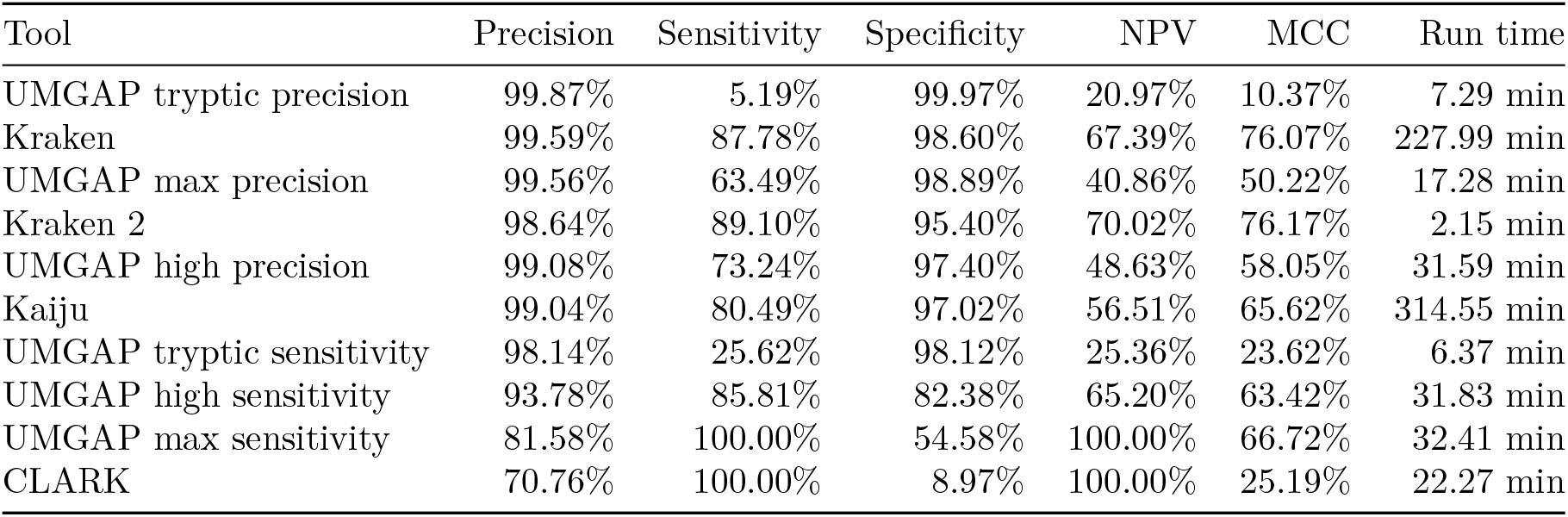
The MetaBenchmark performance metrics for ten metagenomics analysis tools at genus level.

**Table 3:** The MetaBenchmark performance metrics for ten metagenomics analysis tools at phylum level.

| Tool | Precision | Sensitivity | Specificity | NPV | MCC | Run time |
| --- | --- | --- | --- | --- | --- | --- |
| UMGAP tryptic precision | 100.00% | 6.10% | 100.00% | 21.03% | 11.32% | 7.27 min |
| Kraken | 99.98% | 88.95% | 99.92% | 69.35% | 78.49% | 223.16 min |
| UMGAP max precision | 99.99% | 73.50% | 99.96% | 48.55% | 59.71% | 17.08 min |
| Kraken 2 | 99.99% | 91.08% | 99.97% | 73.70% | 81.91% | 2.14 min |
| UMGAP high precision | 99.96% | 83.02% | 99.88% | 59.56% | 70.25% | 31.74 min |
| Kaiju | 99.96% | 84.32% | 99.88% | 61.47% | 71.92% | 317.04 min |
| UMGAP tryptic sensitivity | 99.69% | 30.77% | 99.62% | 26.55% | 28.24% | 6.40 min |
| UMGAP high sensitivity | 94.72% | 93.35% | 83.48% | 79.83% | 75.68% | 31.85 min |
| UMGAP max sensitivity | 81.50% | 100.00% | 37.88% | 100.00% | 55.56% | 32.52 min |
| CLARK | 74.28% | 100.00% | 7.87% | 100.00% | 24.17% | 22.39 min |

**Figure 10:**
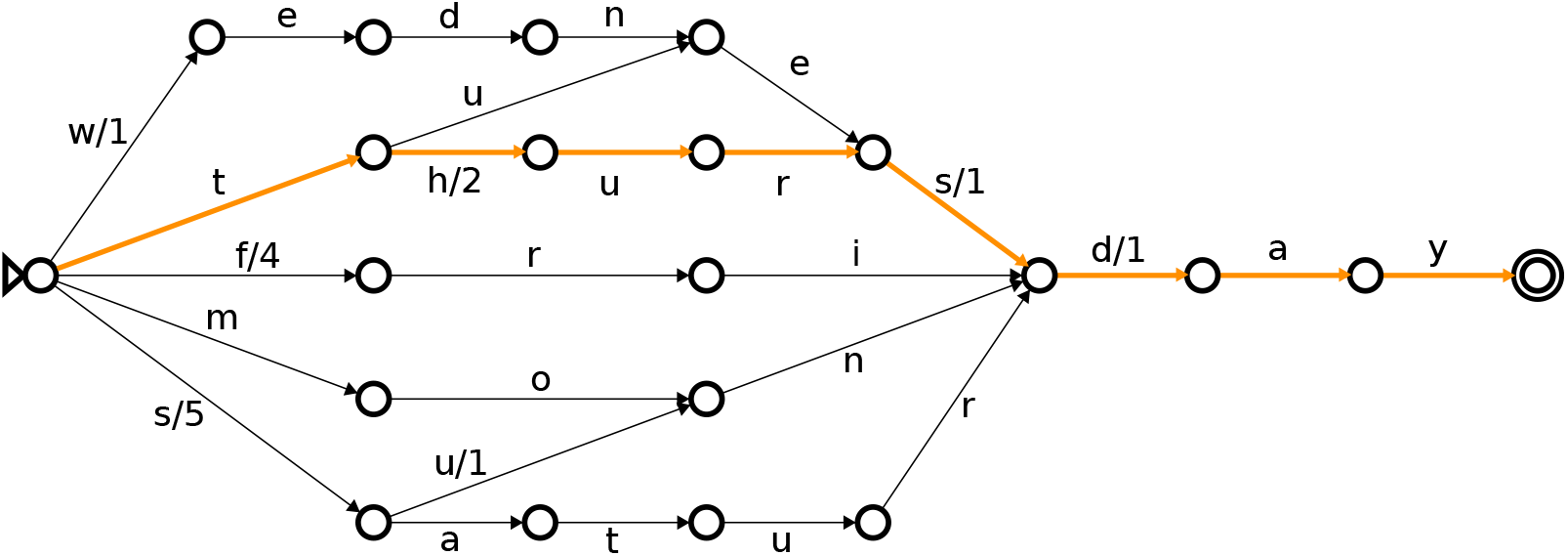
Finite state transducer mapping all weekdays to their index number (monday = 1, tuesday = 2, …). Integer labels are not shown on edges with zero weight. Adding weights along the path spelled by the letters of the word Thursday, from the initial state on the left (indicated by a triangle) to the final state on the right (indicated by a double circle), yields 2 + 1 + 1 = 4. So, Thursday is the fourth day in the week.

**Figure 11:**
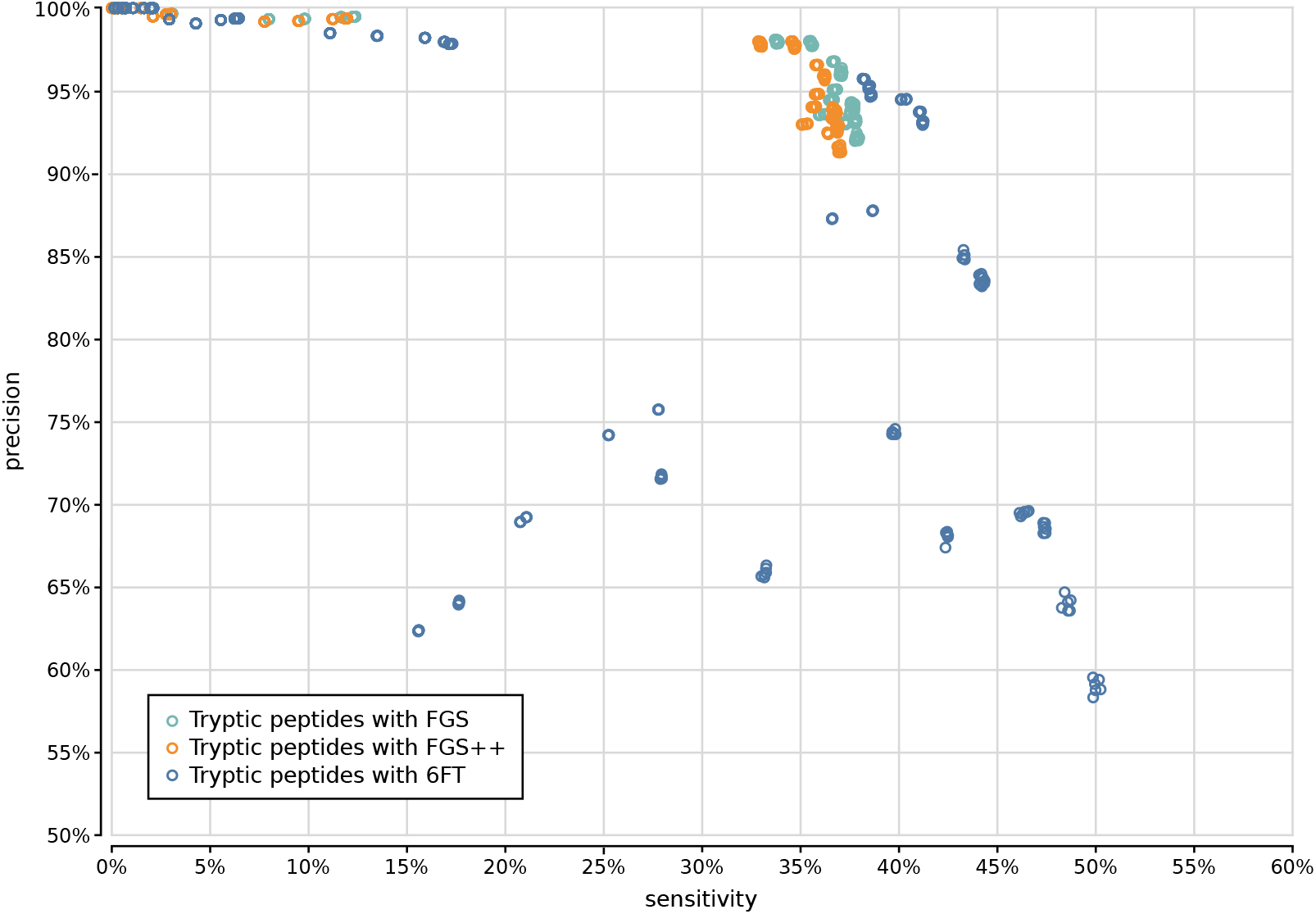
Precision and sensitivity of 2700 tryptic UMGAP configurations, classified per protein translation method: configurations using FGS gene predictions marked in cyan, configurations using FGS++ gene predictions marked in orange and configurations using six-frame translations (6FT) marked in blue.

**Figure 12:**
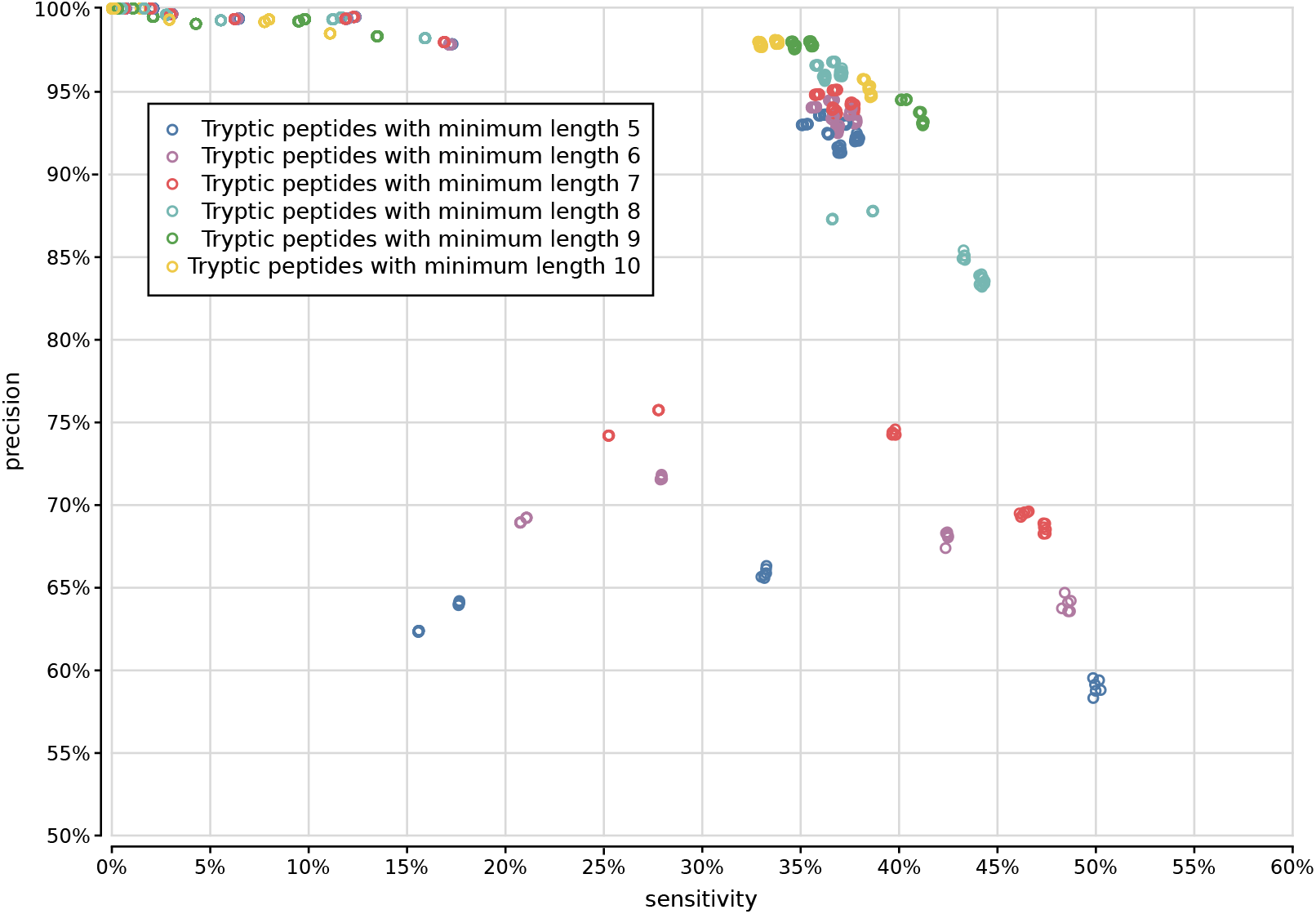
Precision and sensitivity of 2700 tryptic UMGAP configurations, classified per minimal peptide length.

**Figure 13:**
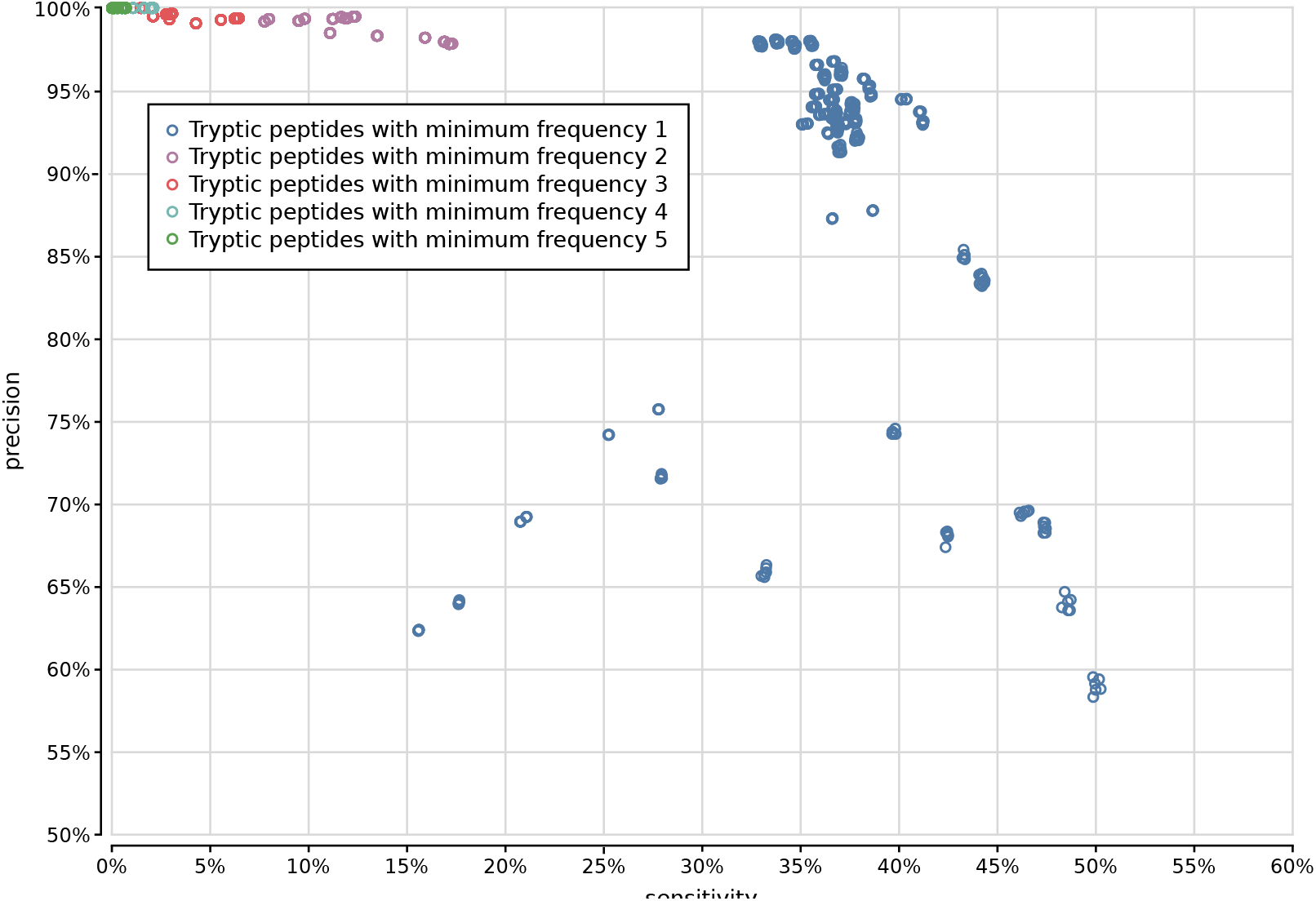
Precision and sensitivity of 2700 tryptic UMGAP configurations, classified according to filtering of low-frequency identifications.

**Figure 14:**
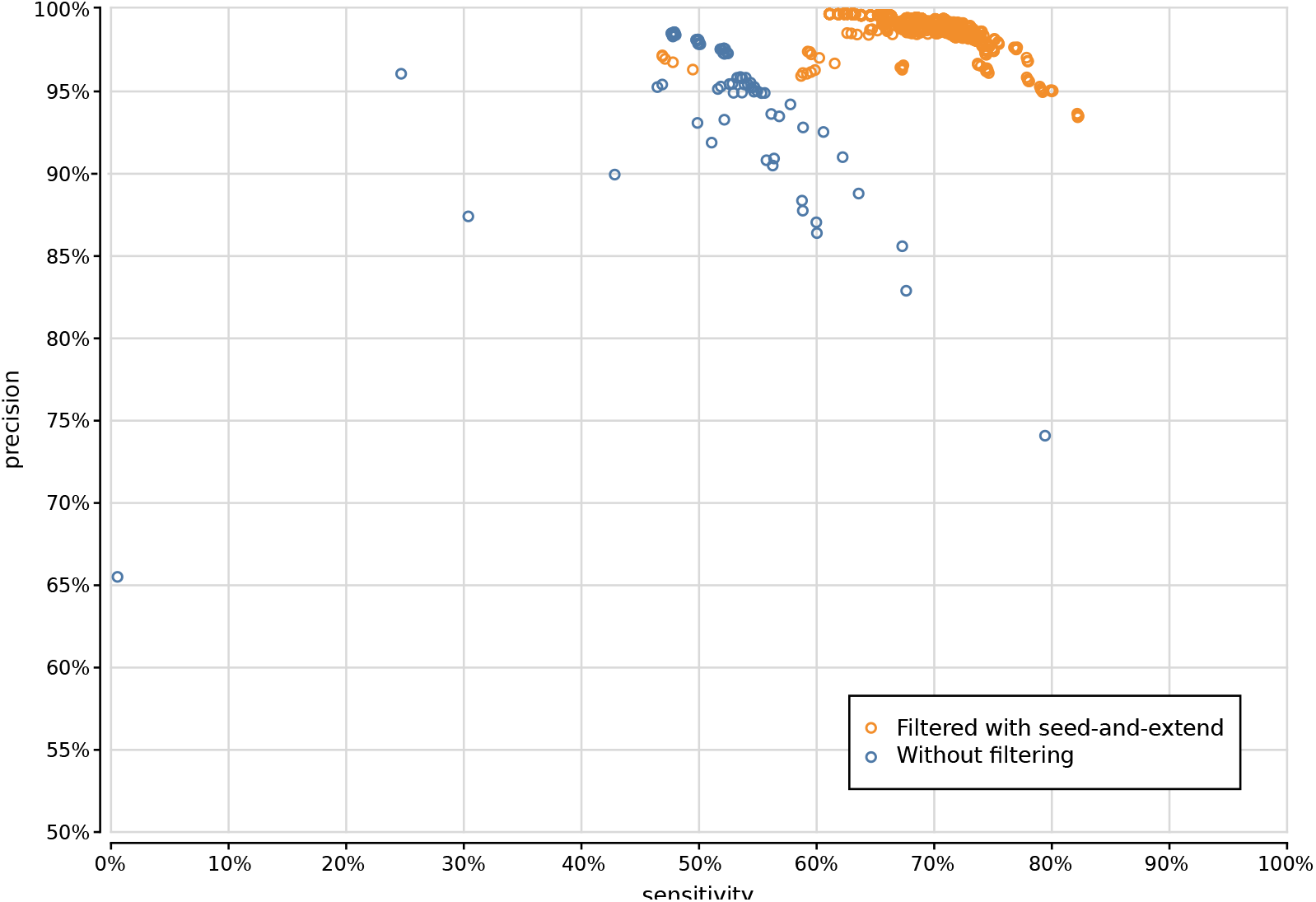
Precision and sensitivity of 1200 UMGAP configurations using 9-mer peptide fragmentation, with configurations that don’t use seed-and-extend filtering marked in blue and configurations that do use seed-and-extend filtering marked in orange.

**Figure 15:**
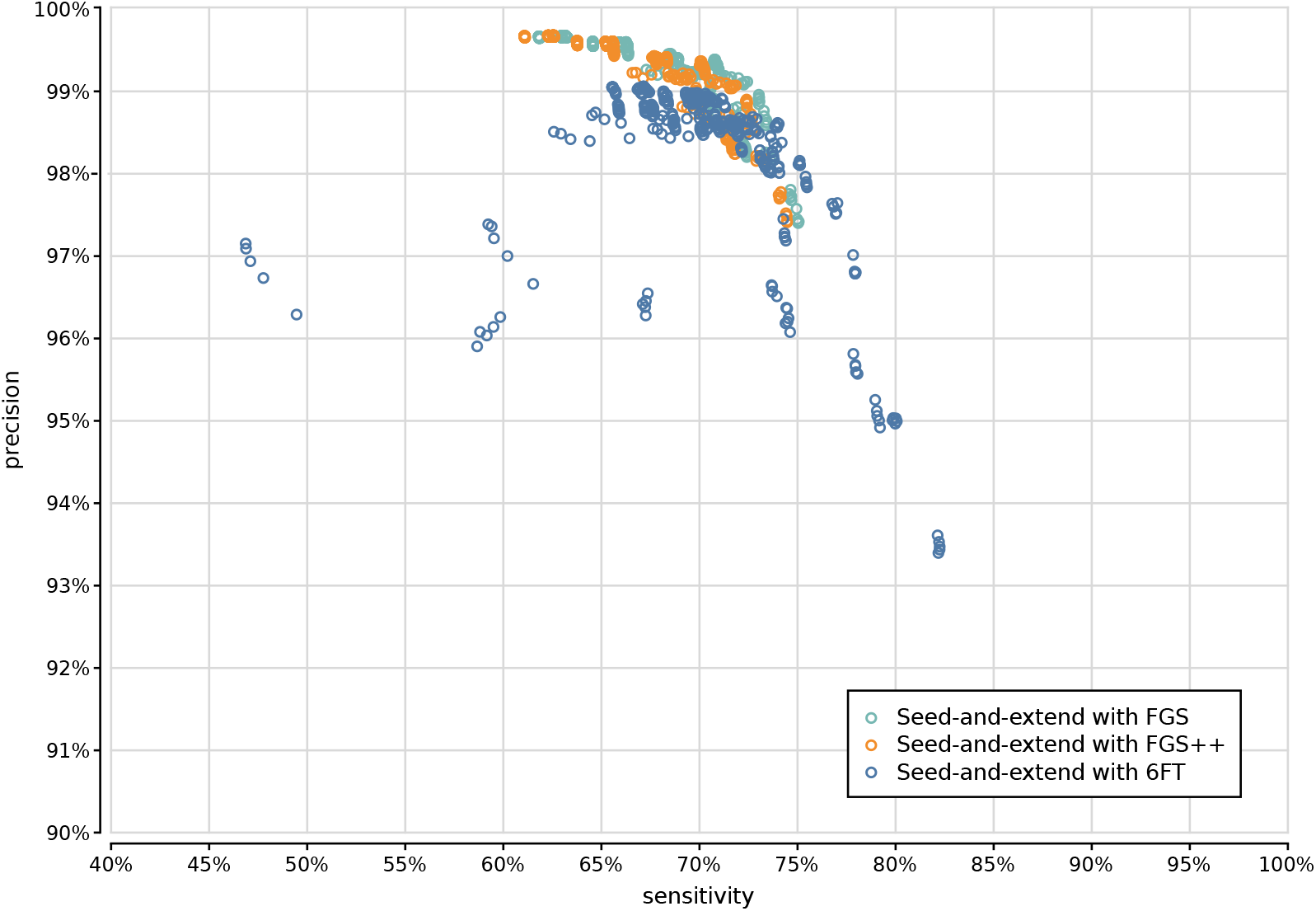
Precision and sensitivity of 1125 UMGAP configurations using 9-mer peptide fragmentation in combination with seed-and-extend filtering, classified per protein translation method: configurations using FGS gene predictions marked in cyan, configurations using FGS++ gene predictions marked in orange and configurations using six-frame translations (6FT) marked in blue.

**Figure 16:**
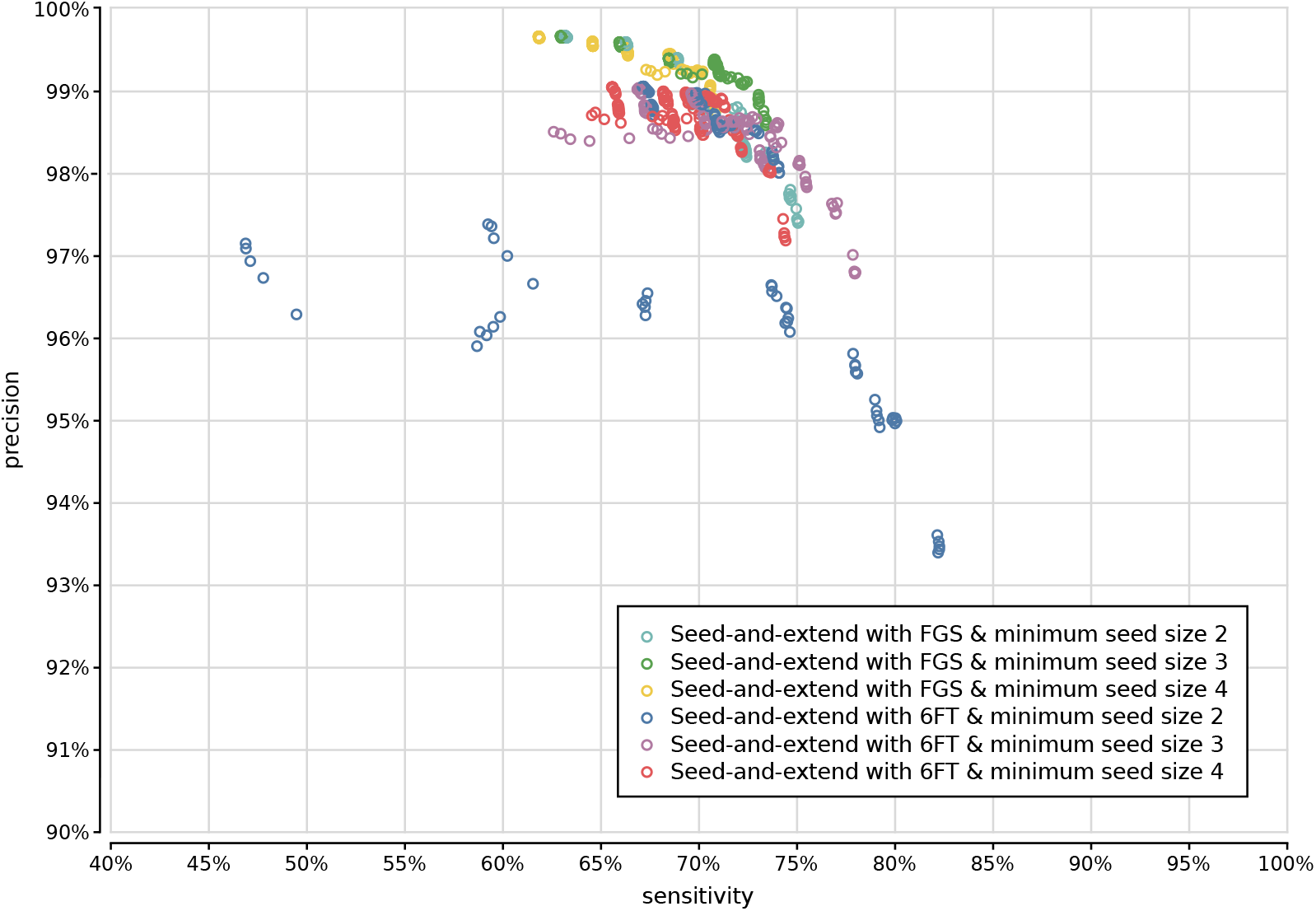
Precision and sensitivity of 750 UMGAP configurations using 9-mer peptide fragmentation in combination with seed-and-extend filtering, classified per minimal seed size and protein translation method. Gene prediction with FGS++ has been excluded.

**Figure 17:**
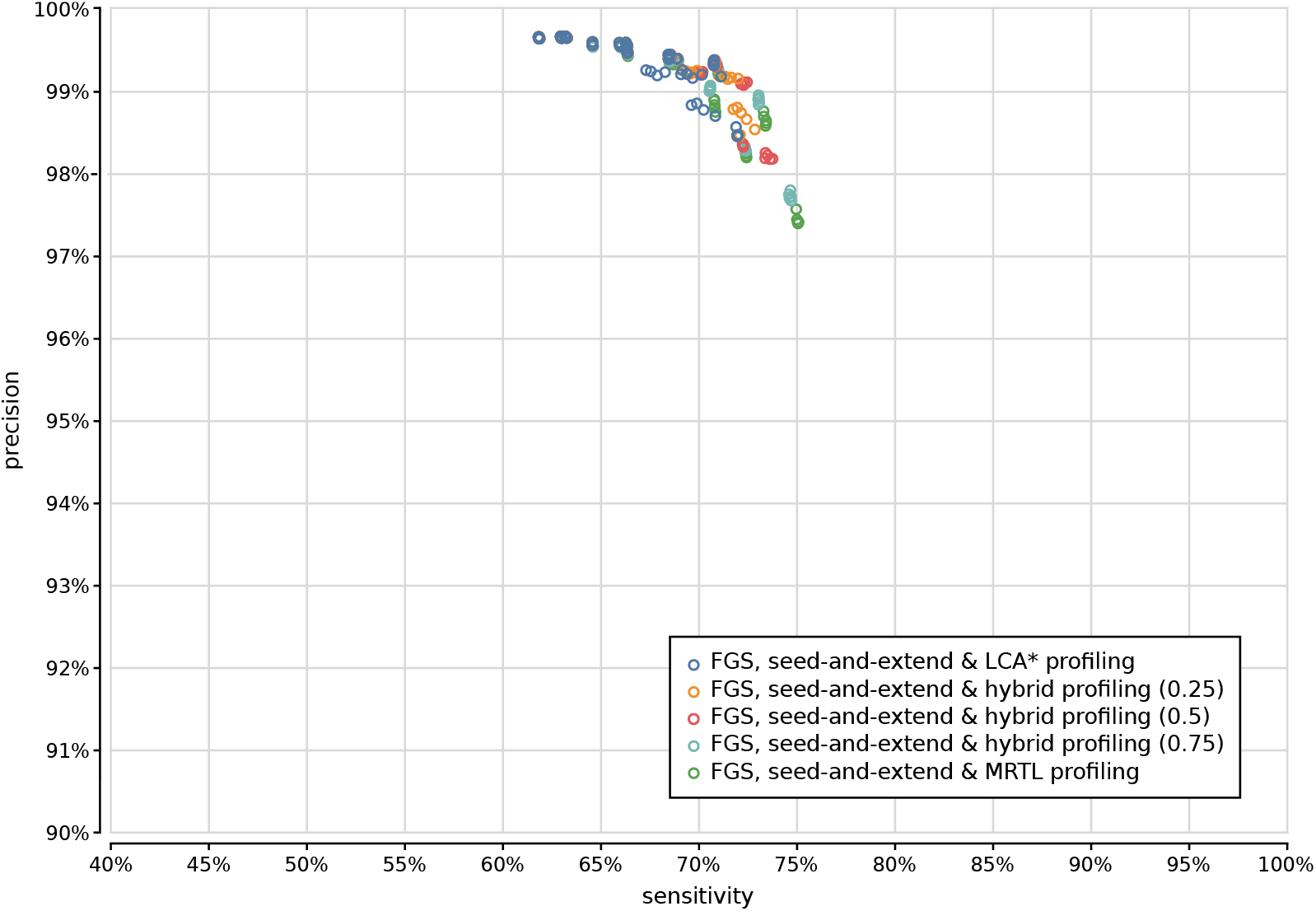
Precision and sensitivity of 375 UMGAP configurations using 9-mer peptide fragmentation, seed-and-extend filtering and gene prediction, classified per read profiling method.

**Figure 18:**
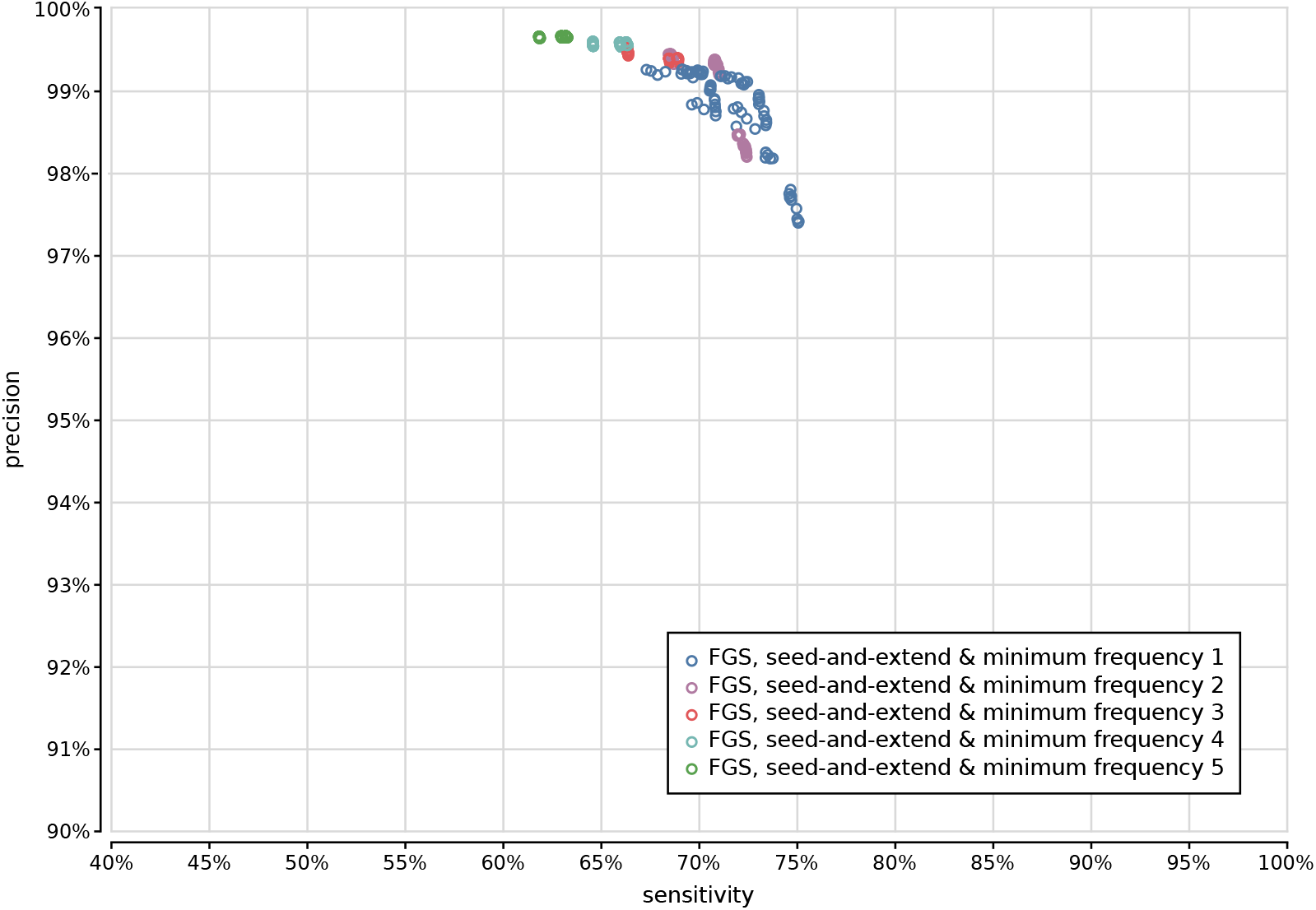
Precision and sensitivity of 375 UMGAP configurations using 9-mer peptide fragmentation, seed-and-extend filtering and gene prediction, classified per filtering of low-frequency identifications.

**Figure 19:**
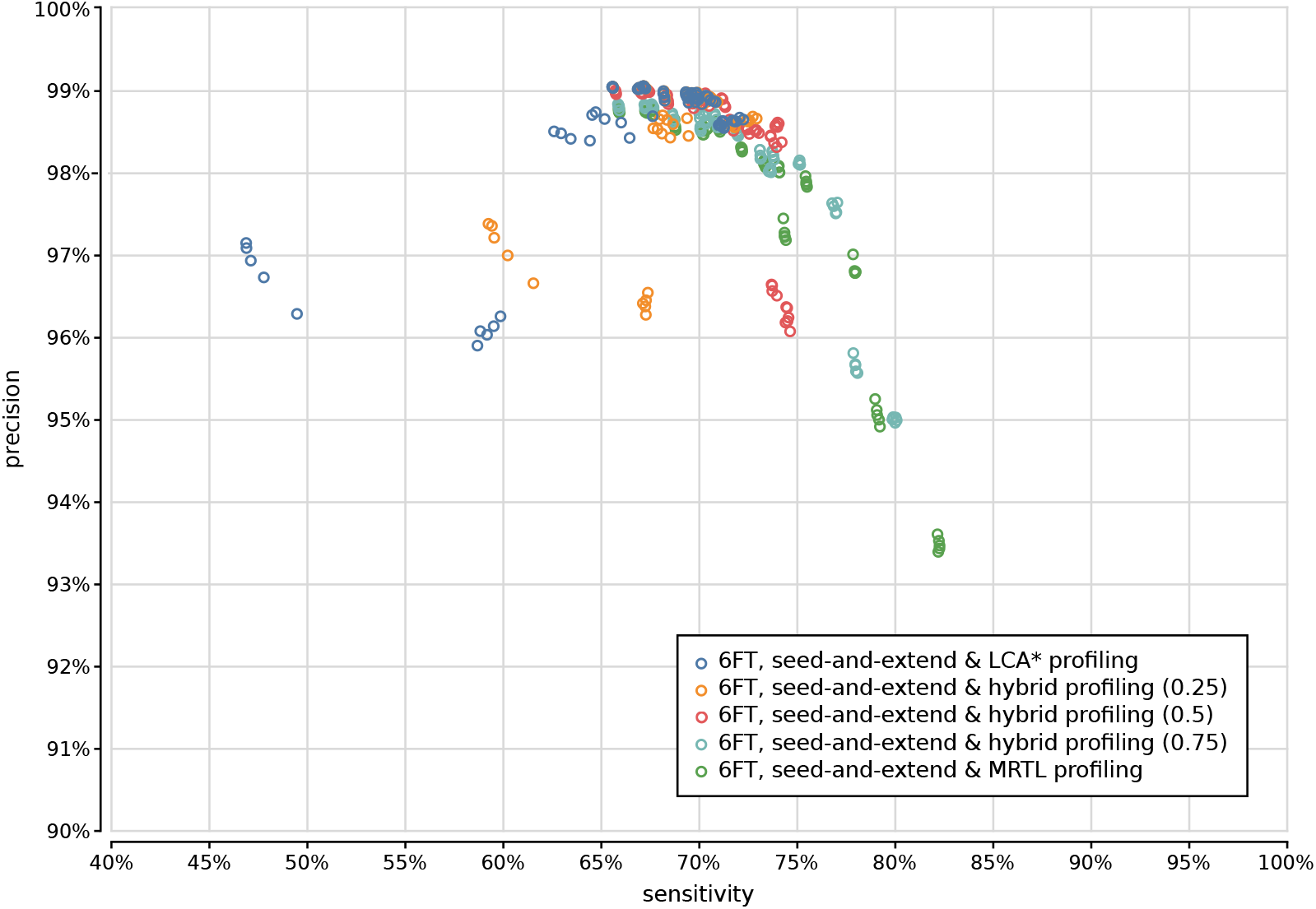
Precision and sensitivity of 375 UMGAP configurations using 9-mer peptide fragmentation, seed-and-extend filtering and six-frame translation, classified per read profiling method.

**Figure 20:**
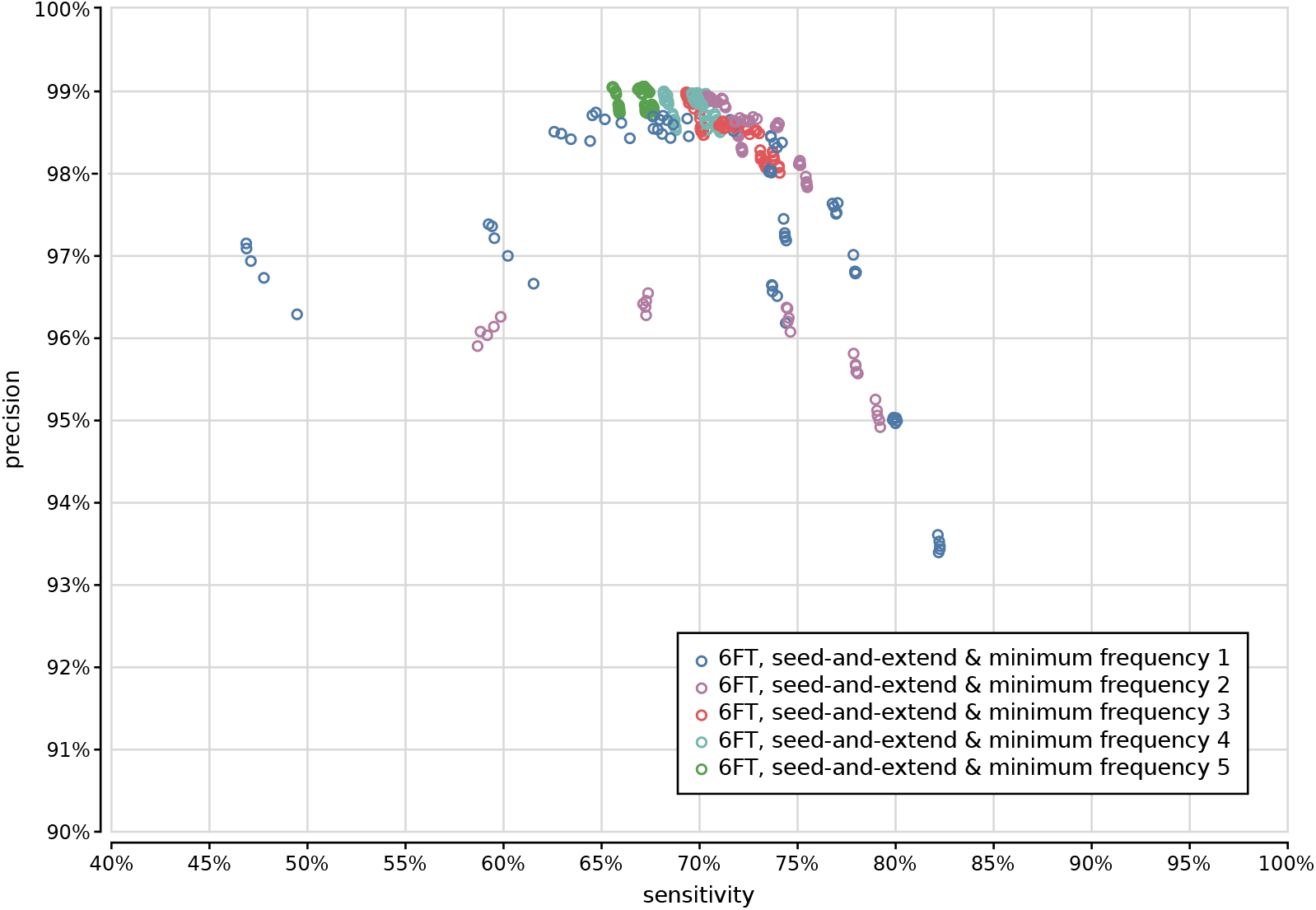
Precision and sensitivity of 375 UMGAP configurations using 9-mer peptide fragmentation, seed-and-extend filtering and six-frame translation, classified per filtering of low-frequency identifications.

## References

Altschul, S.F., Gish, W., Miller, W., Myers, E.W., and Lipman, D.J. 1990. “Basic Local Alignment Search Tool.” Journal of Molecular Biology 215 (3): 403–10.

Boeckmann, B., A. Bairoch, R. Apweiler, M.C. Blatter, A. Estreicher, E. Gasteiger, M.J. Martin, et al. 2003. “The SWISS-PROT Protein Knowledgebase and Its Supplement TrEMBL in 2003.” Nucleic Acids Res. 203 (31): 365–70.

Boisvert, S., Raymond, F., Godzaridis, E., Laviolette, F., and Corbeil, J. 2012. “Ray Meta: Scalable de Novo Metagenome Assembly and Profiling.” Genome Biol. 13: 1–13.

Brady, A., and Salzberg, S.L. 2009. “Phymm and PhymmBL: Metagenomic Phylogenetic Classification with Interpolated Markov Models.” Nature Methods 6: 673–76.

Federhen, S. 2012. “The NCBI Taxonomy Database.” Nucleic Acids Research 40 (database issue): 136–43.

Fischer, J., and Heun, V. 2011. “Space-Efficient Preprocessing Schemes for Range Minimum Queries on Static Arrays.” SIAM Journal on Computing 40: 465–92.

Gallant. n.d. “Index 1,600,000,000 Keys with Automata and Rust.” Accessed November 25, 2016. https://blog.burntsushi.net/transducers/.

Gurdeep Singh, R., Tanca, A., Palomba, A., Van der Jeugt, F., Verschaffelt, P., Uzzau, S., Martens, L., Dawyndt, P., and Mesuere, B. 2019. “Unipept 4.0: Functional Analysis of Metaproteome Data.” Journal of Proteome Research 18 (2): 606–15.

Hugenholtz, P., and G. W. Tyson. 2008. “Metagenomics.” Nature 455: 481–83.

Huson, D.H., Mitra, S., Ruscheweyh, H.J., Weber, N., and Schuster, S.C. 2011. “Integrative Analysis of Environmental Sequences Using Megan4.” Genome Research 21 (9): 1552–60.

Kim, D.J., Hahn, A.S., Wu, S.J., Hanson, N.W., Konwar, K.M., and Hallam, S.J. n.d. “FragGeneScan-Plus for Scalable High-Throughput Short-Read Open Reading Frame Prediction.” In.

Lindgreen, S., Adair, K.L., and Gardner, P.P. 2016. “An Evaluation of the Accuracy and Speed of Metagenome Analysis Tools.” Sci Rep. 6: 19233.

Magrane, M., and UniProt Consortium. 2011. “UniProt Knowledgebase: A Hub of Integrated Protein Data.” Database (Oxford) 2011: bar009.

Menzel, P., Ng, K.L., and Krogh, A. 2016. “Fast and Sensitive Taxonomic Classification for Metagenomics with Kaiju.” Nature Communications 7: 11257.

Mesuere, B., Debyser, G., Aerts, M., Devreese, B., Vandamme, P., and Dawyndt, P. 2015. “The Unipept Metaproteomics Analysis Pipeline.” Proteomics 15 (8): 1437–42.

Mesuere, B., Devreese, B., Debyser, G., Aerts, M., Vandamme, P., and Dawyndt, P. 2012. “Unipept: Tryptic Peptide-Based Biodiversity Analysis of Metaproteome Samples.” J. Proteome Res. 11 (12): 5773–80.

Namiki, T., Hachiya, T., Tanaka, H., and Sakakibara, Y. 2012. “MetaVelvet: An Extension of Velvet Assembler to de Novo Metagenome Assembly from Short Sequence Reads.” Nucleic Acids Res. 40 (20): e155.

Ounit, R., Wanamaker, S., Close, T.J., and Lonardi, S. 2015. “CLARK: Fast and Accurate Classification of Metagenomic and Genomic Sequences Using Discriminative k-Mers.” BMC Genomics 16.

Pell, J., Hintze, A., Canino-Koning, R., Howe, A., Tiedje, J.M., and Brown, C.T. 2012. “Scaling Metagenome Sequence Assembly with Probabilistic de Bruijn Graphs.” Proc. Natl. Acad. Sci. U. S. A. 109 (33): 13272–77.

Peng, Y., Leung, H.C.M., Yiu, S.M., and Chin, F.Y.L. 2011. “Meta-IDBA: A de Novo Assembler for Metagenomic Data.” Bioinformatics 27 (13): i94–101.

Peng, Y., Leung, H.C.M., Yiu, S.M., and Chin, F.Y.L. 2012. “IDBA-UD: A de Novo Assembler for Single-Cell and Metagenomic Sequencing Data with Highly Uneven Depth.” Bioinformatics 28 (11): 1420–28.

Quince, C., Walker, A.W., Simpson, J.T., Loman, N.J., and Segata, N. 2017. “Shotgun Metagenomics, from Sampling to Analysis.” Nature Biotechnology 35: 833–44.

Raymond, E. S. 2003. The Art of UNIX Programming. USA: Addison-Wesley Professional.

Rho, M., Tang, H., and Ye, Y. 2010. “FragGeneScan: Predicting Genes in Short and Error-Prone Reads.” Nucleic Acids Res. 38 (20): e191.

Simpson, J.T., Wong, K., and Jackman, S.D. 2009. “ABySS: A Parallel Assembler for Short Read Sequence Data.” Genome Research 19: 1117–23.

Vandermarliere, E., Mueller, M., and Martens, L. 2013. “Getting Intimate with Trypsin, the Leading Protease in Proteomics.” Mass Spectrom. Rev. 32 (6): 453–65.

Verschaffelt, P., Van Thienen, P., Van Den Bossche, T., Van der Jeugt, F., De Tender, C., Martens, L., Dawyndt, P., and Mesuere, B. 2020. “Unipept CLI 2.0: Adding Support for Visualizations and Functional Annotations.” Bioinformatics 36 (14): 4220–21.

Watson, J.D., Baker, T.A., Bell, S.P., Gann, A., Levine, M., and Losick, R. 2008. Molecular Biology of the Gene. USA: Pearson/Benjamin Cummings.

Wood, D.E., Lu, J., and Langmead, B. 2019. “Improved Metagenomic Analysis with Kraken 2.” Genome Biol. 20 (1): 257.

Wood, D.E., and Salzberg, S.L. 2014. “Kraken: Ultrafast Metagenomic Sequence Classification Using Exact Alignments.” Genome Biol. 15 (3): R46.

